# Antigen receptor signaling and cell death resistance controls intestinal humoral response zonation

**DOI:** 10.1101/2023.03.16.532481

**Authors:** Fiona Raso, Shuozhi Liu, Mikala JoAnn Willett, Gregory Barton, Christian Thomas Mayer, Oliver Bannard, Mridu Acharya, Jagan R. Muppidi, Ann Marshak-Rothstein, Andrea Reboldi

## Abstract

Immunoglobulin A (IgA), the main antibody isotype found in the intestine, has evolved to maintain the stability of commensal communities, and prevent dysbiosis. In stark contrast to systemic antibody response against pathogens, the generation of IgA against intestinal resident microbes assures the simultaneous binding to multiple and diverse commensal-derived antigens. However, the exact mechanisms by which B cells mount such broadly reactive IgA response to the gut microbiome at the mucosal barrier remain elusive. Here we show surface IgA B cell receptor (BCR) is required to confer enhanced B cell fitness during the germinal center reaction in Peyer’s patches and to mediate selection of gut-homing plasma cells with higher efficiency. We demonstrate that, upon antigen stimulation, IgA+ BCR drives greater intracellular signaling in mouse and human B cells and as consequence, IgA+ B cells received higher positive selection cues in the germinal center. Mechanistically, *in vivo* IgA BCR signaling offsets Fas-mediated cell death to rescue low affinity B cell clones and redirects the humoral response to an increased variety of commensal strains at the intestinal interface. Our findings revealed a new mechanism linking tissue-specific antigen receptor signaling with B cell fate and localization of antibody production; and have implications for understanding how intestinal antigen recognition shapes humoral immunity in health and disease.

## INTRODUCTION

Secretion of Immunoglobulin A (IgA) at the intestinal interface is critical to control the ecology of commensal communities: data from both animal models and human studies indicate that deficiency in IgA responses alters the composition of the microbiome^1–3^, leading to increased susceptibility to enteric pathogens and reduced tissue responses to oral vaccines.^4,5^

In recent years, the specificity of IgA has been the focus of intense investigation: it is widely accepted that a single IgA can bind multiple unrelated bacterial taxa^6–9^ a property described as cross-species reactivity ^10^, but species-specific IgA also exist.^11^ While IgA can coat commensals in the absence of T cells ^12–14^, there is evidence that IgA selection is regulated by T cells.^15^ Furthermore, IgA-secreting plasma cells in the intestine are highly mutated^16–20^ and require affinity maturation to properly control commensals.^21^ Somatic mutations and affinity maturation occur in the germinal center (GC) reaction and depend on T cell help, which suggests the GC may play an important role in establishing a functional intestinal IgA response. Peyer’s patches (PPs), the secondary lymphoid organs on the anti-mesenteric side of the small intestine^22^, are characterized by ongoing germinal centers (GC)^23^, and seminal work has identified PPs as the main site for the generation of IgA against intestinal antigens.^24,25^ In PPs, tissue-resident cells produce environmental cues that enforce IgA class switch recombination before B cell entry into the GC^26–28^, and control differentiation of antigen-specific plasma cells.^29^ The GC reaction is primarily required for the selection of B cell clones binding antigen with high affinity, that eventually differentiate into antibody secreting plasma cells^30–32^, a feature which is difficult to reconcile with the known cross-species reactivity of IgA. However, in contrast to peripheral lymph nodes or spleen, GC regulation in PPs has been shown to be peculiar— less dependent on positive and negative T follicular helper signals.^33,34^ Moreover, recent data have highlighted that the complexity of gut microbiome shapes both affinity and clonality during the GC reaction: under homeostatic conditions, selection of highly dominant B cell clones is rare in PPs, a strikingly different situation compared to peripheral lymph nodes upon immunization.^30,35,36^ Thus, a PP-specific mechanism must exist that modulates the GC reaction and underpins the generation of poly/cross-species reactive intestinal IgA.

Following acquisition of AID-mediated BCR point mutations in the dark zone, GC B cells test their mutated BCR for binding of antigen displayed by FDCs and receive T cell help by cognate T follicular helper cells (Tfh). The relative contribution of T cell help and BCR signaling in controlling GC selection has been debated.^37–41^, with the current view is that Tfh are central in discriminating between B cells carrying BCR with different affinities for the antigen and drive differentiation of high affinity B cells into plasma cells.^32^ Nevertheless, several groups have reported that surface BCR isotypes can intrinsically modulate B cell differentiation, including selection during the germinal center.^42–45^

The impact of the IgA BCR on B cells during the response to intestinal commensals remains undefined. The role of IgA BCR in the GC reaction of Peyer’s patches, and its effect on the humoral immune response at the intestinal barrier, remain largely unknown.

In the present study, we examined the role of IgA in regulating chronic GC reactions that form in response to commensal and pathogenic bacteria. We found that IgA facilitates GC B cell fitness and is required for the generation of gut-homing plasma cells. IgA+ B cells are positively selected in the light zone and are resistant to apoptosis: the IgA BCR mediates stronger intracellular signaling and offsets FasL-dependent apoptosis. In conclusion, acquisition of an IgA BCR in PPs is not only required for the generation of low inflammatory antibodies that can be translocated into the lumen, but it also lowers the threshold of selection to ensure a comprehensive response to commensals.

## RESULTS

### IgA+ B cells dominate the germinal center reaction and are preferentially selected into the memory compartment

We initially sought to evaluate whether surface IgA BCR was required for effective participation in the GC reaction. To this end, we used mice with targeted deletion of the IgA switch region and sequence, which completely abolishes both IgA-expressing B cells and secreted IgA.^46^ Immunohistochemical analysis of PPs did not reveal altered GC structure in the absence of IgA (**Figure 1A**), with normal size of the follicle and GC (**Figure S1A)**. We then analyzed PP and mesenteric lymph node (mLN) B cell populations by flow cytometry for the distribution of follicular B cells (B220+ IgD+ GL7-), GC B cells (B220+ IgD- GL7+), memory B cells (B220+ IgD- GL7- CD38+ CD138-), and plasma cells (B220+/- IgD- GL7- CD138+) (**Figure S1B**). No difference in B cell subsets was observed in both PPs (**Figures 1C-E, S1C**) and mLN (**Figure S1D**) between mice deficient for IgA (*Iga*^*-/-*^) and co-housed, littermate controls (*Iga*^*+/+*^), despite an altered distribution of antibody (Ab) isotypes (**Figures S1E-J**).

**Figure 1.**
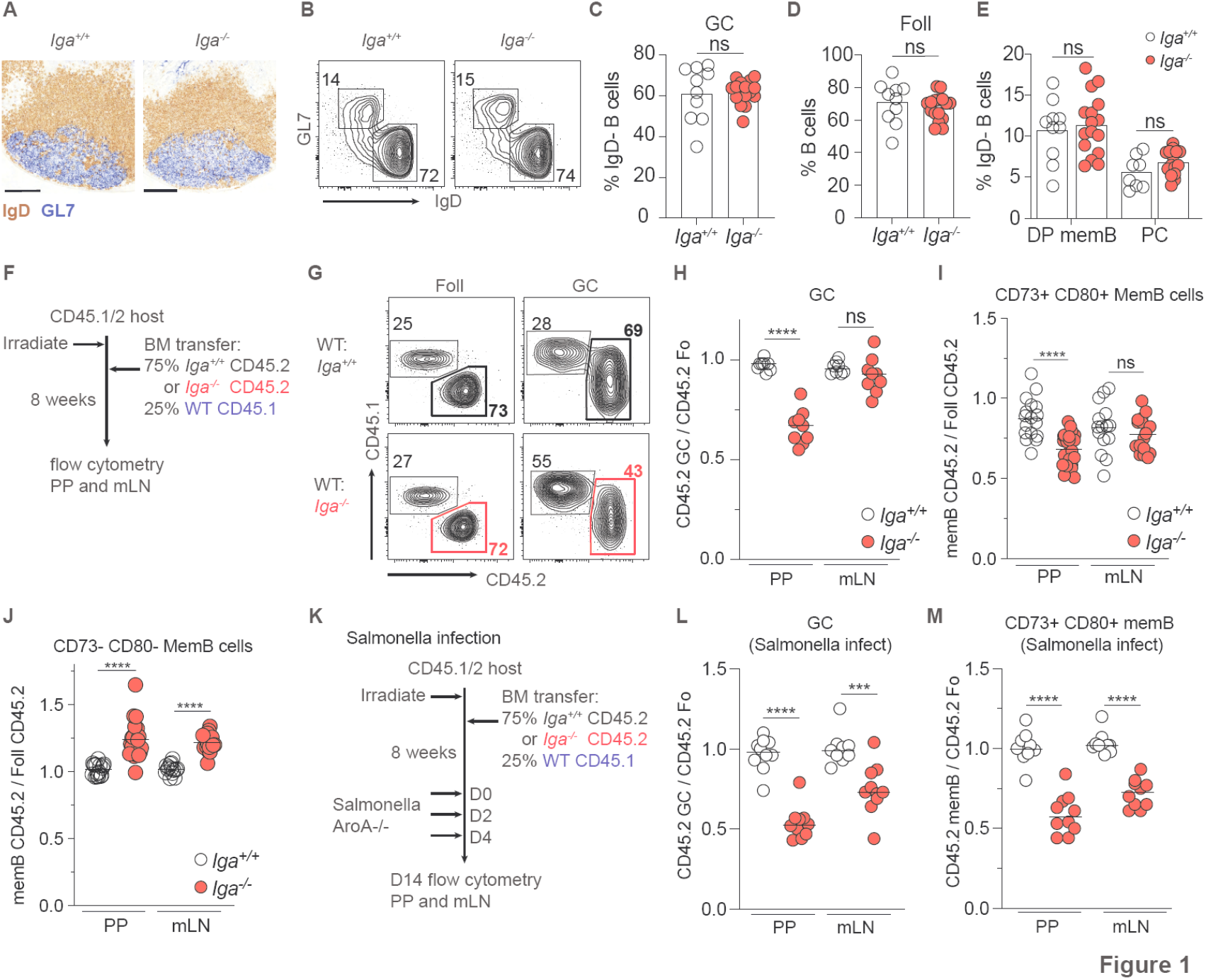
IgA+ B cells dominate the germinal center reaction and are preferentially selected into the memory compartment. (A – E) Steady state, co-housed, 8-12 week old *Iga*^*+/+*^ and *Iga*^*-/-*^ mice. (A) Frozen sections of PPs from *Iga*^*+/+*^ and *Iga*^*-/-*^ mice stained with immunohistochemistry (IHC) antibodies against IgD (brown) and GL7 (blue). Scale bars, 200 um. (B) Representative flow cytometry of PP GC (GL7+ IgD-) and Foll (GL7-IgD+) B cells in *Iga*^*+/+*^ and *Iga*^*-/-*^ mice. Gated previously on live B220+ CD138-B cells. (C) PP GC B cells as percentage of activated B cells. (D) PP Foll as percentage of B cells. (E) CD73+ CD80+ (DP) MemB, PCs as percentage of activated (IgD-) B cells. A-E, compiled data from at least 3 independent experiments with 1-3 mice per group. (F) *Iga*^*-/-*^ mBM chimera experimental set up for G - J. (G) Representative flow cytometry of CD45 staining on Foll and GC B cells in mBM chimeras. (H) Ratio of frequency of CD45.2 GC B cells to CD45.2 Foll B cells in PP and mLN. (I) Ratio of frequency of CD45.2 DP memB cells to CD45.2 Foll B cells in PP and mLN. (J) Ratio of frequency of CD45.2 DN memB cells to CD45.2 Foll B cells in PP and mLN. (K) *Iga*^*-/-*^ mBM chimera and Salmonella infection experimental set up for L - M. (L) Ratio of frequency of CD45.2 GC B cells to CD45.2 Foll B cells in PP and mLN of Salmonella infected chimeras. (M) Ratio of frequency of CD45.2 DP memB cells to CD45.2 Foll B cells in PP and mLN of Salmonella infected chimeras. Unpaired t-test was used in c, d, e, h, i, j, l, m. ns=not significant, *p<0.005, **p<0.001,***p<0.0005, ****p<0.0001.

However, while PP B cells expressing isotypes other than IgA can effectively become GC B cells, suggesting that IgA is not required for GC formation, it does not rule out a role for the IgA BCR during the GC reaction. GC selection is an iterative immune process controlled by competition among B cell clones^47^, thus, we wanted to test whether IgA was required in a competitive setting. To this end, we generated mixed bone-marrow (BM) chimeric mice where most B cells (75%) were from *Iga*^*-/-*^ or *Iga*^*+/+*^ CD45.2+ donor and the remaining B cells (25%) were from wild-type (WT) CD45.1+ donor (**Figure 1F**). Flow cytometry analysis revealed that in PPs, *Iga*^*-/-*^ B cells were largely impaired in their ability to participate in the GC reaction compared to WT B cells, while follicular and GC B cells were present at the same ratio in the control mixed BM chimeras (**Figures 1G, 1H**). No defect was observed in mLN of either mixed BM chimeras, likely due to the low frequency of IgA+ B cells participating in the GC reaction and their peculiar TGFβ-independent^27^, commensal-independent^36^ nature. The inability of *Iga*^*-/-*^ B cells to compete in the GC was also apparent in immunohistochemistry staining of PP frozen sections, with CD45.2 *Iga*^*-/-*^ B cells filling the B cell follicle, while inefficiently entering the GC structure, which was instead dominated by CD45.1 WT B cells. This pattern was reversed in control mixed BM, with CD45.2 *Iga*^*+/+*^ B cells being more abundant in both B cell follicle and GC (**Figure S1K**). GC B cells can give rise to both plasma cells and memory B cells. Memory B cell expressing CD73 and CD80 are thought to be directly generated in GCs^48^. In line with reduced participation in the GC reaction, mixed BM chimeras revealed that *Iga*^*-/-*^ B cells also contributed less to GC-derived memory in PPs, but not mLN (**Figure 1I**). In contrast, IgA deficient B cells were overrepresented in the double-negative (CD73-CD80-) compartment (**Figure 1J**), suggesting that IgA is required for optimal GC participation and formation of GC-derived memory.

Chronic GC reactions in PPs and mLN are mainly driven by commensals but can also incorporate responses to pathogens. To test whether intrinsic IgA-deficiency impacts GC B cell competition during infection, we infected mixed BM chimera with metabolically deficient Salmonella strain (ΔAroA) which is able to induce an IgA response (**Figure 1K**).^49^ Similar to homeostatic conditions, *Iga*^*-/-*^ B cells were largely outcompeted in GC of PPs, but not mLN (**Figure 1L**). Interestingly, in contrast to what was observed during commensal driven responses, DP MemB cells were affected by IgA deficiency in both PPs and mLN upon Salmonella infection (**Figure 1M**) while DN MemB were largely unchanged (**Figure S1L**). This discrepancy could be due to enhanced migration of PP-generated DP MemB cells into mLN during Salmonella infection compared to homeostatic situations.

Together our data show that IgA is required for optimal GC persistence and memory B formation during responses to both commensal and pathogenic enteric bacteria.

### IgA is required for efficient plasma cell homing to the intestinal lamina propria

In addition to memory B cells, GC B cells can also differentiate into plasma cells (PCs), and selection during the GC reaction is thought to be critical for the generation of PCs that secrete high affinity Abs. PP GCs generate most IgA-secreting PCs that egress from lymphatics at the basal side of the PPs and home to the lamina propria ^50^ via the integrin alpha 4 beta 7 (α4β7)^51,52^ and C-C chemokine receptor type 9 (CCR9).^53^ Indeed, the vast majority of PCs expressing α4β7 and CCR9 in PPs were IgA+, while PCs that did not express both α4β7 and CCR9 (non-gut homing) were not enriched as IgA expressing cells in PPs (**Figure 2A**).

**Figure 2.**
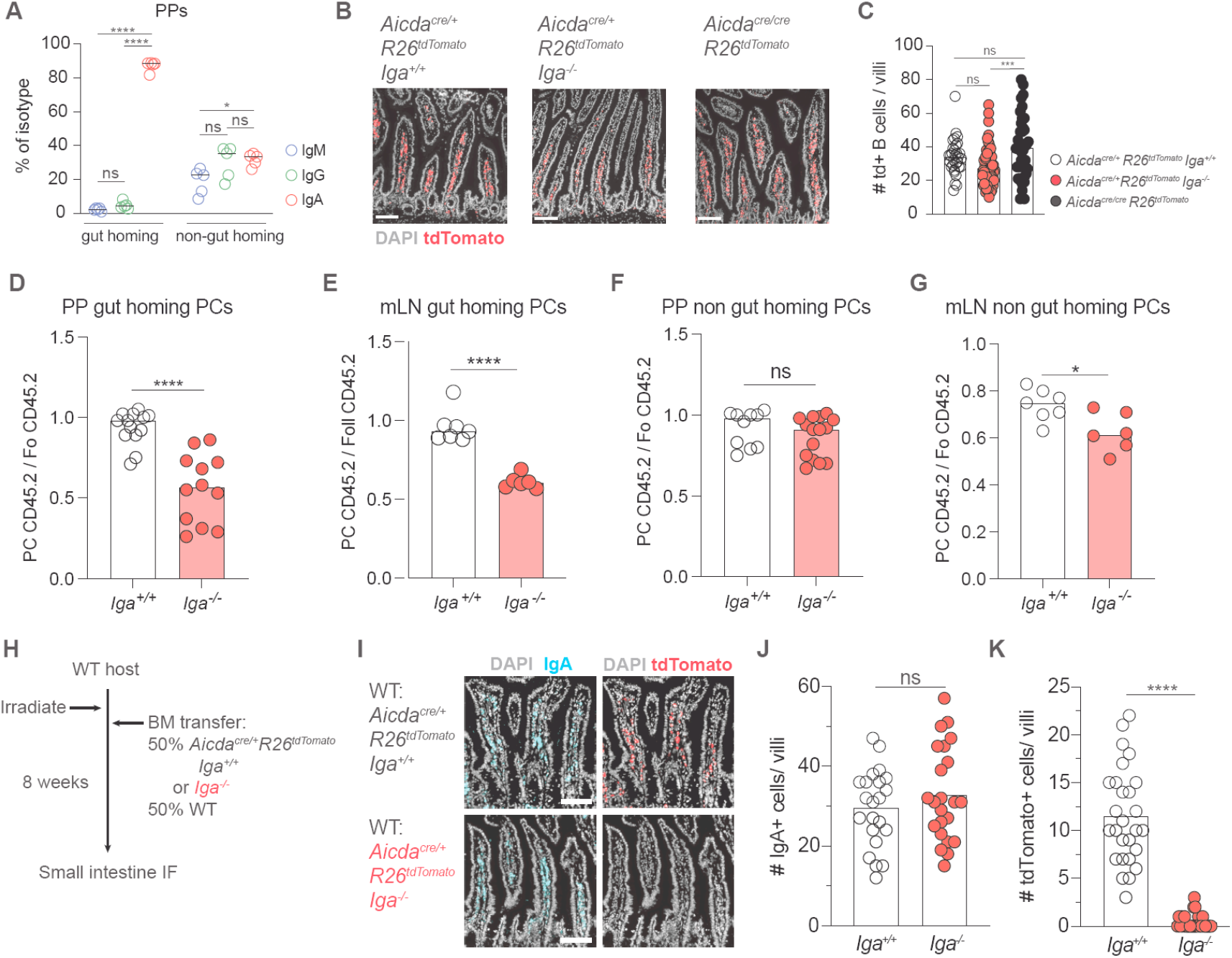
IgA is required for efficient plasma cell homing to the intestinal lamina propria. (A) Isotypes of gut homing (α4β7+ CCR9+) and non-gut homing PCs from PPs of WT mice treated with FTY-720 for seven days. Compiled data from 2 independent experiments with 2-3 mice per group. (B) Representative immunofluorescence of small intestinal villi from *Aidca*^*cre/+*^ R26^tdTomato^ *Iga*^*+/+*^, *Aidca*^*cre/+*^ R26^tdTomato^ *Iga*^*+/+*^, *Aidca*^*cre/cre*^ R26^tdTomato^ mice stained with DAPI (gray) and tdTomato (red). Scale bars, 100 um. (C) Quantification of multiple images as in (B). Number of tdTomato+ cells per villi. Each symbol represents one villi with at least 5 villi quantified per image. Data from 2-3 mice per group. (D) Ratio of frequency of CD45.2 gut homing PCs (defined as CCR9+ α4β7+ CD138+) to CD45.2 Foll B cells from PPs of mBM chimeras. (E) Ratio of frequency of CD45.2 non-gut homing PCs to CD45.2 Foll B cells from PPs of mBM chimeras. (F) Ratio of frequency of CD45.2 gut homing PCs (defined as CCR9+ α4β7+ CD138+) to CD45.2 Foll B cells from mLN of mBM chimeras. (G) Ratio of frequency of CD45.2 non-gut homing PCs to CD45.2 Foll B cells from mLN of mBM chimeras. Data from D-G are representative of at least 3 independent experiments with 3-5 mice per group. (H) *Aidca*^*cre/+*^ R26^tdTomato^ *Iga*^*-/-*^ mBM chimera experimental set up for I - K. (I) Representative immunofluorescence of small intestinal villi stained for DAPI (gray), IgA (cyan), and tdTomato (red) in WT: *Aidca*^*cre*^ R26^tdTomato^ *Iga*^*+/+*^ or *Iga*^*-/-*^ mBM chimeras. Scale bars, 100 um. (J, K) Quantification of multiple images as in I. (J) Number of IgA+ cells per villi. (K) Number of tdTomato+ cells per villi. Data from I-K are representative of 2 independent experiments with 2-3 mice per group. Each symbol represents one villi with at least 5 villi quantified per image. One way anova was used in a and c. Unpaired t-test was used in d, e, f, g, j, k. ns=not significant, *p<0.005, **p<0.001,***p<0.0005, ****p<0.0001.

Although IgA+ GC B cells make up a sizable share of GC B cells (**Figure S1E**), they were present at a lower frequency compared to IgA+ PCs (**Figure S1H**), suggesting an IgA-dependent mechanism for selection of gut-homing PCs. PP-generated PCs traffic through the mLN on their way to reach the lamina propria: similar to PPs, most α4β7 and CCR9 expressing PCs were IgA+ (**Figure S2A)**. Interestingly, absence of IgA does not impair gut homing, as microscopy analysis of fluorescent PCs in the intestinal lamina propria of mice unable to switch to IgA (*Aidca*^cre/+^ Rosa26^Stop-tdTomato/+^ *Iga-/-* and *Aidca*^cre/cre^ Rosa26^Stop-tdTomato/+^) did not show decreased PC compartment compared to IgA-sufficient mice (*Aidca*^cre/+^ Rosa26^Stop-tdTomato/+^ *Iga*^*+/+*^) (**Figures 2B, 2C, S2B**).

However, PC generation from GC B cells is dependent on selection. In mixed chimeras while the total *Iga*^*-/-*^ PCs were slightly reduced (**Figure S2C**), a marked reduction was observed in gut-homing *Iga*^*-/-*^ PCs in both PPs (**Figure 2D**) and mLN (**Figure 2E**) while non-gut homing PCs were minimally altered (**Figure 2F, 2G**). To determine whether *Iga*^*-/-*^ PCs were nonetheless able to reach the lamina propria, we generated mixed BM chimeras in which half of PCs, either *Iga*^*+/+*^ or *Iga*^*-/-*^, were fluorescently marked (*Aidca*^cre/+^ Rosa26^Stop-tdTomato/+^ *Iga*^*+/+*^ or *Iga*^*-/-*^) (**Figure 2H**). While in the *Iga*^*+/+*^: WT chimera the intestine was repleted with IgA+ PCs, from either WT (non-fluorescent) or *Iga*^*+/+*^ B cells (tdTom+) (**Figures 2I, 2J, S2D**), *Iga*^*-/-*^ PCs (tdTom+) were not observed in the intestine of *Iga*^*-/-*^: WT chimera, suggesting that IgA-deficiency impairs homing to the LP in a competitive setting (**Figures 2I, 2K**).

Taken together, these data point to a critical role of surface IgA to instruct the generation of gut-homing plasma cells. It appears that, while non-IgA expressing B cells can differentiate into PCs, they are largely unable to upregulate homing receptors required for efficient migration into the intestinal lamina propria.

Mixed BM chimeras are a robust experimental system to investigate competitive requirements for immunological processes. However, the need for irradiation can introduce alterations due to cell death, especially in the gut ^54^. Therefore, to further expand our findings to a competitive system that does not require irradiation, we took advantage of allelic exclusion, a process by which B cells only transcribe one heavy chain locus during development. As a result of this, IgA heterozygous (*Iga*^*+/-*^) mice normally develop two genetically distinct sets of B cells, one sufficient for IgA and one deficient for IgA, effectively generating a natural WT:*Iga*^*-/-*^ chimera (**Figure 3A**). Since WT and *Iga*^*-/-*^ cells carry distinct immunoglobulin heavy chain allotypes, allotypic reagents can be used to distinguish B cells from WT (IgH^b^) and *Iga*^*-/-*^ (IgH^a^) origin. Due to allelic exclusion, follicular B cells are present at a 50:50 ratio in both WT (IgH^a/b^) and *Iga*^*+/-*^ (IgH^a/b^): if surface IgA does not preferentially enhance GC fitness, we expect to observe a 50% reduction of total IgA GC B cells in *Iga*^*+/-*^ mice compared to WT mice, due to the *Iga*^*-/-*^ B cells equal contribution to the GC. However, if surface IgA promotes competitiveness in the GC, we anticipate the overall IgA class switch to be unaffected in *Iga*^*+/-*^ mice. Indeed, we found that total IgA GC B cells were indistinguishable between WT and *Iga*^*+/-*^ mice, supporting the concept of IgA dominance during GC competition (**Figure 3B**). In contrast, IgM^a^ B cells (derived from *Iga*^*-/-*^ BM) were expanded in activated B cell subsets compared to IgM^b^ WT B cells (**Figures 3C, S2E**), a finding also observed in mixed BM chimeras (**Figure S2F**), suggesting that intrinsic IgA deficiency during GC competition allows the persistence of IgM+ GC B cells. Allotypic reagents can be used to track the origin of intestinal Ab: flow cytometry analysis of Ab-coated intestinal bacteria^7,55^ revealed that the mucosal humoral response was dominated by IgA+ cells (**Figures 2D, S2G**) and unaffected in *Iga*^*+/-*^ mice, a finding in accordance with the requirement of IgA for intestinal plasma cell generation (**Figures 2I, 2K)**. While no changes were observed in commensal-reactive serum IgA between WT and *Iga*^*+/-*^ mice (**Figure S2H**), *Iga*^*+/-*^ PCs showed increased contribution to commensal-reactive IgM in serum (**Figures 2E, S2I**), suggesting that IgA-deficient B cells are able to respond to commensals and differentiate into PCs, but their response is redirected into systemic circulation. To assess whether a similar mechanism was in place during an antigen-specific response to foreign antigen, we orally immunized *Iga*^*+/-*^ and WT mice with cholera toxin (CT)^56^ (**Figure 2F**). Analysis of Abs from in vitro differentiated memory B cells revealed the anti-CT Ab was dominated by IgA from *Iga*^*+/+*^ B cells (**Figure 2G**), while *Iga*^*-/-*^ B cells increasingly generated anti-CT IgM Abs (**Figures 2H, I**). Together our data support a model where surface IgA BCR allows for efficient participation in the PP GC and PC generation, underpinning efficient humoral immune responses at the intestinal barrier.

**Figure 3.**
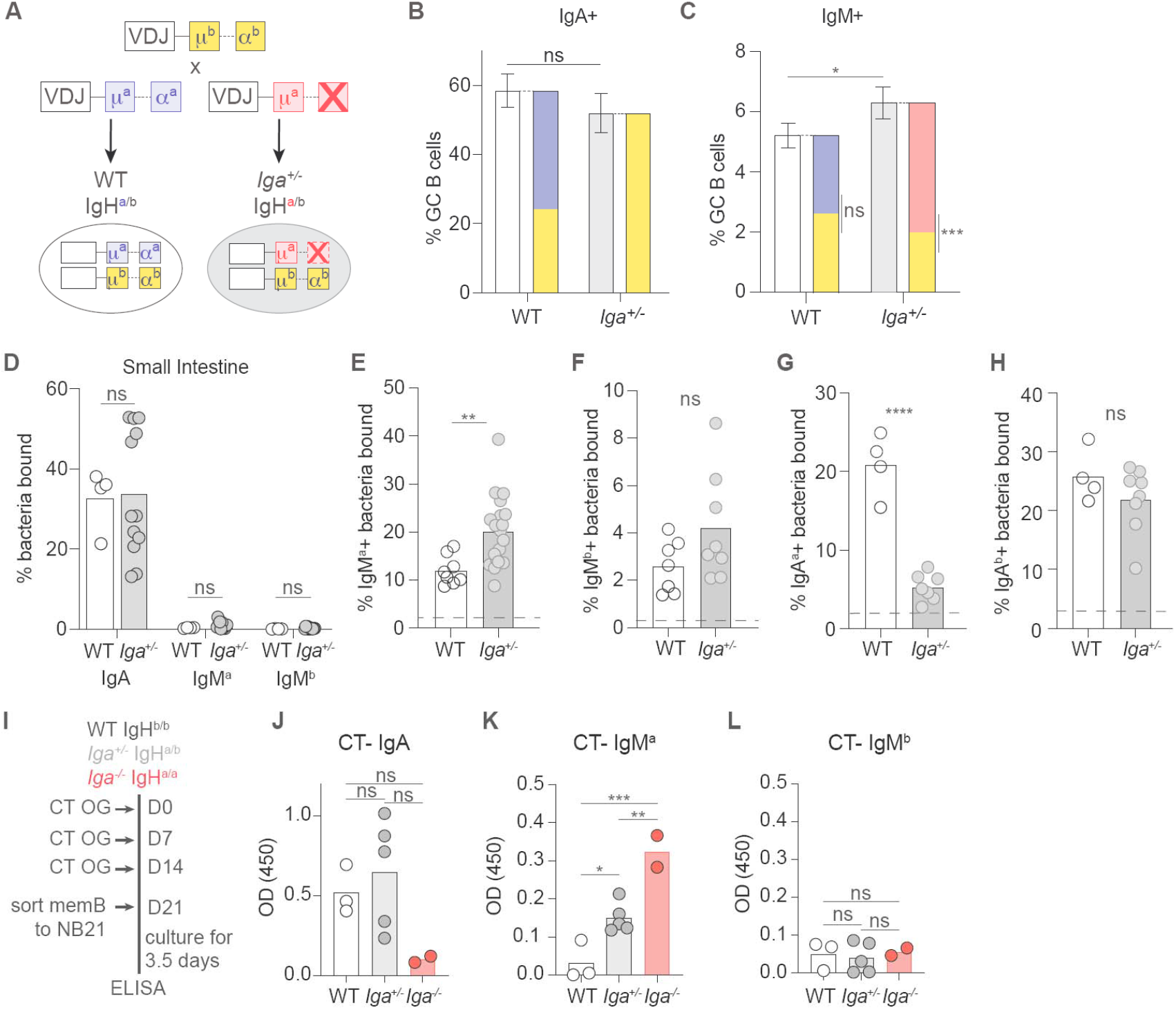
IgA is dominant in activated B cells of intrinsic chimera mice. (A) Breeding setup for allelic exclusion in WT IgH^a/b^ and *Iga*^*+/-*^ IgH^a/b^ mice. (B, C) Percentage of IgA+(B) and IgM+ (C) GC B cells in PPs. Solid bars represent indicated total isotype in GC; stacked bars represent allotypic isotype in GC of WT and *Iga*^*+/-*^ mice. (D) Percentage of endogenously coated small intestinal bacteria with indicated antibodies. (E, F) Percentage of intestinal commensal from uMT mice stained by IgM^a^ (E) or IgM^b^ (F) antibodies from *Iga*^*+/+*^ or *Iga*^*+/-*^ serum. (G, H) Percentage of intestinal commensal from uMT mice stained by IgA^a^ (G) or IgA^b^ (H) antibodies from *Iga*^*+/+*^ or *Iga*^*+/-*^ serum. Dashed line represents unstained uMT bacteria in E-H. (D – H) Complied data from at least 3 independent experiments with 2-3 mice per group. (I) Experimental setup for cholera toxin (CT) oral immunization and subsequent memory B cell in vitro culture on NB21 feeder cells. (J, K, L) ELISA OD for CT specific IgA (J), IgM^a^ (K), and IgM^b^ (L) antibodies detected in memory B supernatant 3.5 days after in vitro culture. J-L, compiled data from 2 independent experiments with 1-3 mice per group. Unpaired t-test was used in b, c, d, e, f, g, h. One-way anova was used in j, k, l. ns=not significant, *p<0.005, **p<0.001,***p<0.0005, ****p<0.0001.

### Resistance to cell death and T cell help enhance IgA+ GC B cell persistence

The GC is divided into anatomically restricted dark zone (DZ) and light zone (LZ) compartments. While flow cytometry analysis of mixed BM chimeras did not show a preferential reduction of *Iga*^*-/-*^ B cells in one GC compartment (**Figures 4A, 4B, S3A**), in situ immunofluorescent microscopy revealed that *Iga*^*-/-*^ B cells were predominantly clustered in the LZ (**Figures 4C, 4D, 4E**). The GC reaction couples massive B cell proliferation with carefully controlled cell death to achieve clonal selection. We therefore initially tested the ability of *Iga*^*-/-*^ B cells to undergo proliferation by measuring BRDU incorporation. *Iga*^*-/-*^ and WT B cells proliferated at similar rates in both GC compartments and in the follicle (**Figures 4F, S3B-E**). In the DZ, GC B cells undergo cell death in response to the AID-dependent generation of a non-functional BCR.^57,58^ To test whether *Iga*^*-/-*^ B cells were lost at this negative selection stage, we retrovirally expressed *Bcl2*, an anti-apoptotic gene that can rescue defects in DZ GC survival^58^ in mixed BM chimeras. Ectopic BCL-2 was unable to rescue *Iga*^*-/-*^ B cells in both GC and memory compartments (**Figures 4G, S3F**), suggesting IgA BCR does not confer competitive advantage during DZ selection.

**Figure 4.**
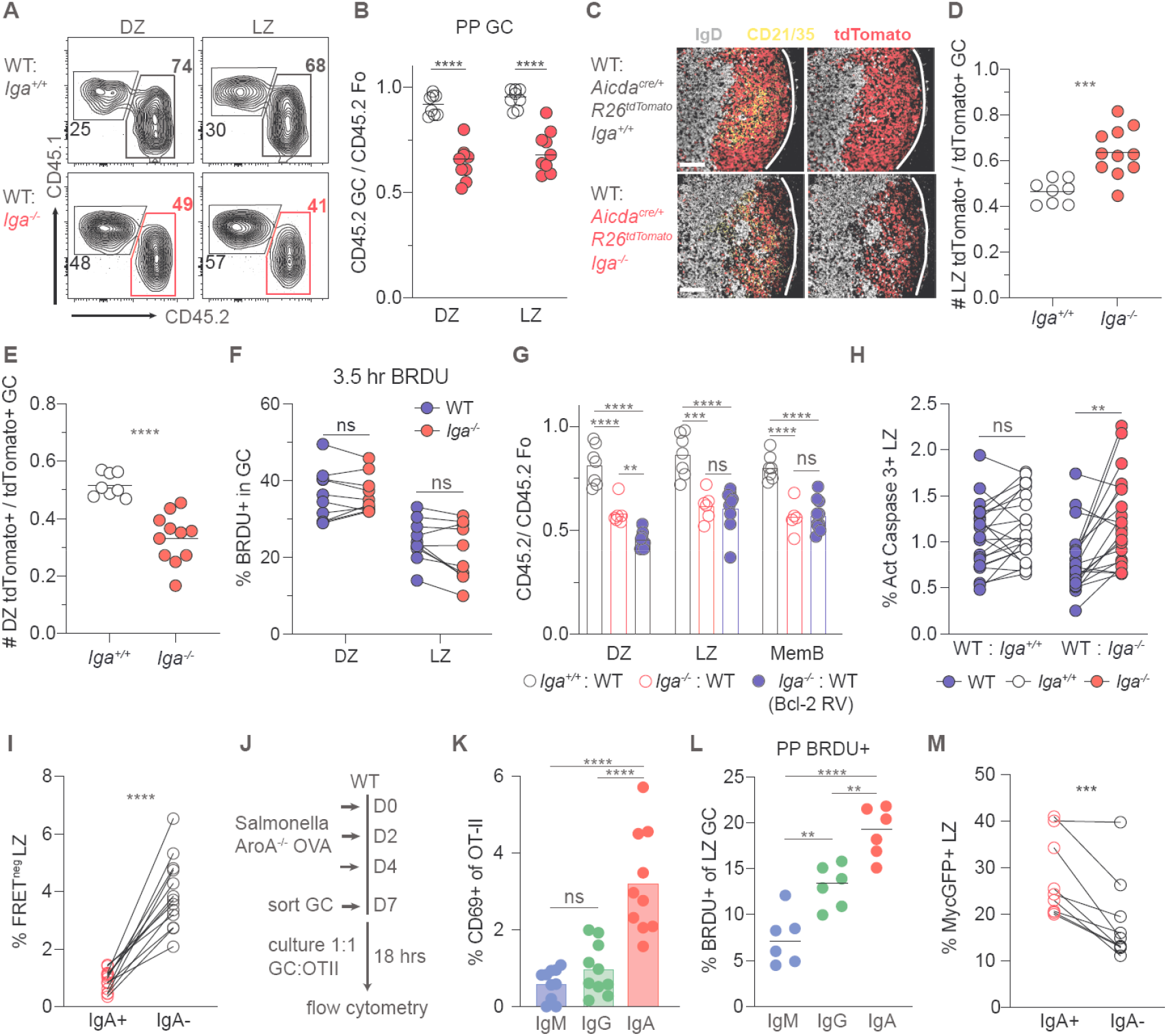
IgA+ B cells exhibit higher cell death resistance and experience stronger T cell-dependent help in GC LZ. (A) Representative flow cytometry of CD45 staining on PP GC LZ and DZ B cells in mBM chimeras. (B) Ratio of frequency of CD45.2 LZ or DZ B cells to CD45.2 Foll B cells in PP of mBM chimeras. Data is representative of at least 3 independent experiments with at least 3 mice per group. (C) Representative immunofluorescence of PPs stained for DAPI (gray), tdTomato (red) and CD21/35 (cyan), in WT: *Aidca*^*cre*^ R26^tdTomato^ *Iga*^*+/+*^ or *Iga*^*-/-*^ mBM chimeras. Scale bars, 100 um. Lines indicate GC edges with LZ oriented on left, DZ right. (D, E) Quantification of multiple images as in **c**. Ratio of frequency of tdTomato+ cells in LZ (D) or DZ (E) to total GC B cells. Data from C-E are representative of 2 independent experiments with 2-3 mice per group. (F) Percentage of BRDU+ DZ or LZ GC B cells in PPs of *Iga*^*-/-*^ :WT mBM chimeras after 3.5 hour BRDU pulse. Compiled data from at least 3 independent experiments with at least 3 mice per group. (G) Ratio of frequency of indicated CD45.2 B cells to CD45.2 Foll B cells in PP of WT BCL2 : *Iga*^*-/-*^ BCL2 retroviral mBM chimeras. Compiled data from 2 experiments with at least 3 mice per group. (H) Percentage of active caspase3+ LZ GC B cells in mBM chimeras. Compiled data from at least 3 independent experiments with at least 3 mice per group. (I) Percentage of FRET negative LZ IgA+ and IgA-cells from *Rosa26*^*INDIA*^ PPs. Compiled data from at least 3 independent experiments with at least 3 mice per group. (J) Experimental set up for Salmonella AroA^-/-^ expressing Ova infection and in vitro GC antigen presentation. (K) Percentage of CD69+ CD4+ OT-II T cells co-cultured with indicated GC B cell isotype. GC subsets were sorted from PPs according to isotype expression (IgG = IgG1 + IgG2b). Each symbol is the average of technical replicates from one mouse. Compiled data from two experiments with 5 mice per group. (L) Percentage of BRDU+ LZ GC B cells in PPs for indicated isotypes after 30-minute BRDU pulse. GC subsets were gated according to isotype expression (IgG = IgM-IgA-). Compiled data from two experiments with 3 mice per group. (M) Percentage of cMycGFP+ LZ IgA+ and IgA-B cells from c-Myc-GFP PPs. Compiled data from at least 3 experiments with at least 3 mice per group. Unpaired t-test was used in b, d, e, f, h, i, m. One-way anova was used in g, k, l. ns=not significant, *p<0.005, **p<0.001,***p<0.0005, ****p<0.0001.

In the LZ, GC B cells are deleted due to the absence of positive selection, which integrates both BCR signaling and T cell help. Cell death analysis in mixed BM chimera by active caspase 3 staining revealed a significantly higher frequency of apoptosis among LZ, but not DZ, *Iga*^*-/-*^ B cells (**Figures 4H, S3G-H**), a phenotype restricted to PPs (**Figure S3I**). Analysis of Rosa26^INDIA^ apoptosis indicator mice which report active caspase 3 via FRET loss^57^ also revealed that IgA BCR confers protection from cell death in the LZ during the GC reaction in PPs (**Figure 4I**). To test whether increased apoptosis resistance in LZ IgA+ B cells was associated with higher degree of T cell help, we orally immunized mice with an attenuated Salmonella strain encoding OVA. The same number of GC B cells were positively sorted according to surface isotypes and co-cultured with naïve OVA-specific TCR transgenic CD4 T cells (**Figure 4J**). IgA+ GC B cells induced higher CD69 expression in CD4 T cells compared to IgM+ or IgG+ GC B cells (**Figure 4K**). Since no difference was observed in MHCII expression (**Figure S3J**), our results suggest that IgA expressing GC B cells are either equipped with a more efficient antigen processing and presentation machinery or are better in extracting antigen from follicular dendritic cells. Recently, it has been shown that LZ T cells metabolically refuel LZ GC B cells without controlling their re-entry into S phase^59^, a process that can be measured by a short pulse of BRDU. We observed that in the PP LZ, IgA+ GC B cells outcompeted other GC isotypes for□ T cell–derived refueling cues as shown by a higher proportion of IgA+ GC B cells had incorporated BRDU (**Figure 4L**), with a similar, albeit more subtle, trend also present in mLN (**Figure S3K**). Correspondingly, IgA+ LZ B cells showed increased c-Myc expression, suggesting integration of stronger positive selection cues (**Figure 4M**). Together our results support the concept that the IgA BCR protects GC B cells from cell death in the LZ via enhanced T cell-derived positive interaction that permits preferential selection during GC response.

### IgA mediates stronger BCR signaling

In the GC LZ, B cell clones that bind antigen with higher affinity are preferentially selected for survival^57^, therefore, we hypothesized that the IgA BCR would mediate stronger BCR signaling. Ex-vivo stimulation of Ca2+ dye-loaded GC B cells with pan anti BCR revealed that, in contrast to IgM+ or IgG2b+ GC B cells which had undetectable Ca2+ responses, as previously reported^37,60^, IgA+ GC B cells spiked calcium readily and consistently (**Figures 5A and 5D**), a feature also observed using isotype-specific Ab stimulation (**Figures 5B and 5E**), despite no difference in surface expression of the BCR complex (**Figure S4A**). To increase sensitivity in BCR-dependent Ca2+ release analysis, we also generated mice that express both the red fluorescent protein Tdtomato and the ultrasensitive calcium indicator GCamp6 in GC and post-GC B cells (*Aidca*^cre/+^ Rosa26^Stop-tdTomato/gCaMP6^) and observed a similar IgA-dependent Ca2+ release (**Figures 5C, 5F, S4B**). In contrast, BCR-dependent Ca2+ spike in memory B cells, albeit faster in IgA+ cells, was observed in all the memory B cell isotypes (**Figures S4C-S4I**), highlighting the difference between GC and memory B cell BCR signaling. The IgA-dependent Ca2+ spike was BCR dependent as incubation with Ibrutinib, a BTK inhibitor, reduced the calcium flux observed in both IgA GC and memory B cells (**Figures 5F, S4J-S4M**) (*22*).

**Figure 5.**
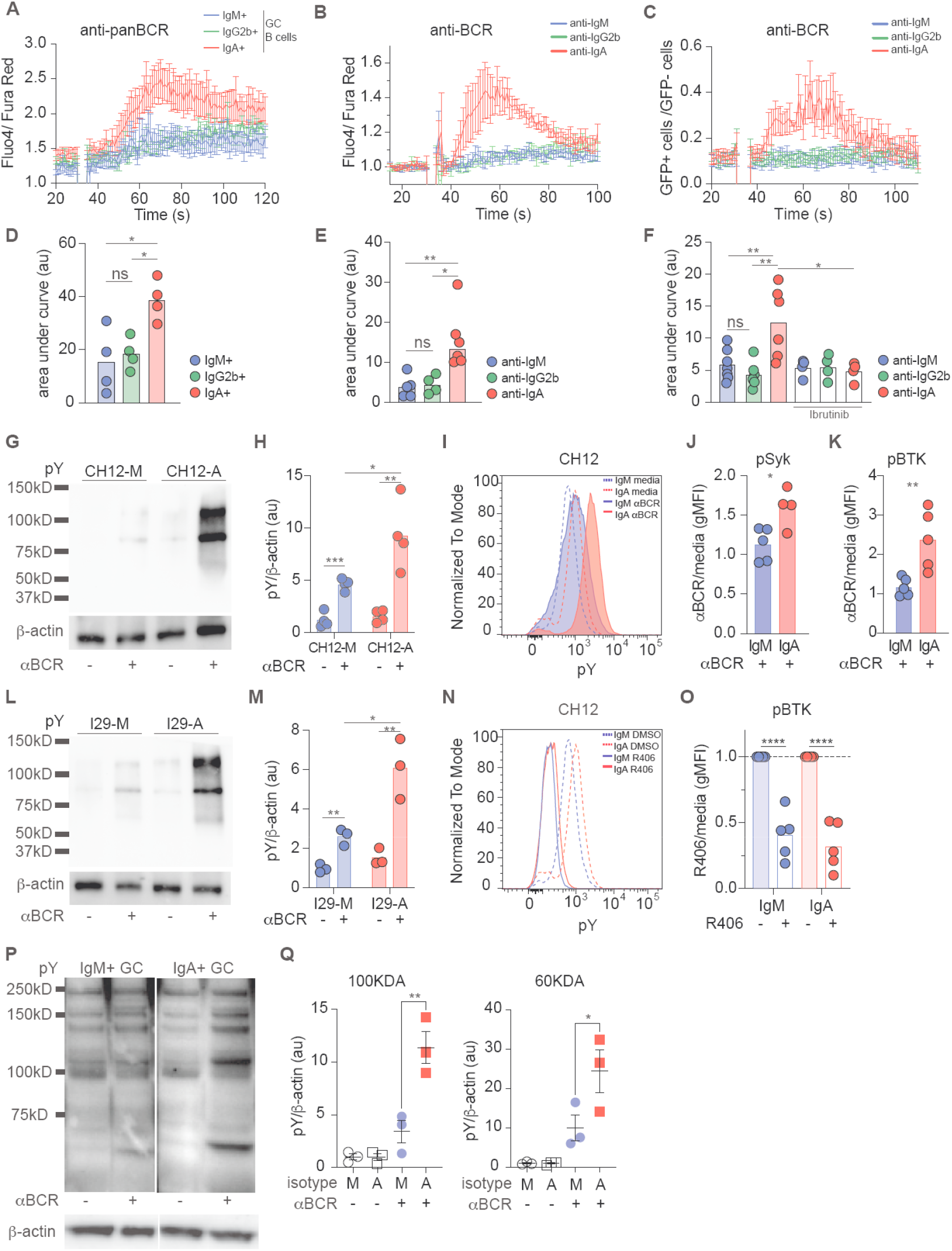
Enhanced BCR signaling in murine and human IgA+ GC B cells. (A) Compiled calcium traces for IgM, IgG2b, and IgA PP GC B cells (defined as isotype negative populations) stimulated with anti-panBCR. (B) Compiled calcium traces for *Aidca*^*cre/+*^ R26^td/+^ PP GC B cells stimulated with indicated anti-BCR isotypes. (C) Compiled calcium traces from *Aidca*^*cre/+*^ R26^td/gCAMP^ PP GC B cells stimulated with indicated anti-BCR isotypes. Shown as ratio of frequency of gCAMP-GFP+ to gCAMP-GFP-cells per second. (D) Area under the curve of individual calcium traces calculated between 50 and 100 seconds for A. A, D are compiled data from 4 independent experiments. (E) Area under the curve of individual calcium traces calculated between 40 and 100 seconds for B. Indicated samples were pre-treated with ibrutinib for 10 minutes prior to anti-BCR stimulation. B, E are compiled data from at least 3 independent experiments. (F) Area under the curve of individual calcium traces calculated between 40 and 100 seconds for **c**. Indicated samples were pre-treated with ibrutinib for 10 minutes prior to anti-BCR stimulation. C, F are compiled data from at least 3 independent experiments. (G) Representative pY western blot for CH12 cells stimulated with anti-BCR. (H) Quantitative analysis from G of pY normalized to β-actin. Compiled data from 4 independent experiments. **i**. Representative histogram of pY staining in CH12 cells after BCR stimulation. (J) pSyk gMFI normalized to untreated in CH12 cells after anti-BCR. (K) pBTK gMFI normalized to untreated in CH12 cells after anti-BCR. (J,K) are compiled data from at least 3 independent experiments. (L) Representative pY western blot for I29 cells stimulated with anti-BCR. (M) Quantitative analysis from L of pY normalized to β-actin. (N) Representative histogram of pY in CH12 cells after incubation with R406. (O) pBTK gMFI normalized to untreated in CH12 cells after incubation with R406. Compiled data from at least 3 independent experiments. (P) Representative pY and β-actin western blots from GC human tonsils of indicated isotypes after anti-BCR stimulation sorted as isotype negative populations. (Q) Quantitative analysis from P of pY normalized to β-actin. Compiled data from 3 independent experiments. Unpaired t-test was used in j, k, o. One-way anova was used in d, e, f, h, m, r. ns=not significant, *p<0.005, **p<0.001,***p<0.0005, ****p<0.0001.

BCR signaling via different isotypes has been shown to mediate various biological outcomes^42,44,45^, but little is known about the impact of the IgA BCR on intracellular signaling. Thus, to investigate IgA BCR signaling with increased granularity, we took advantage of CH12 and I29, two unrelated murine B cell lines with the ability to class switch to IgA^61,62^, to generate CH12 and I29 as IgM (CH12-M; I29-M) or IgA (CH12-A; I29-A) expressing lines (**Figures S5A and S5B**). Anti-isotype stimulation of CH12 revealed a more profound phosphorylation pattern in CH12-A compared to CH12-M by both western blot and phosphoflow (**Figures 5G, 5H, 5I, S5C**), including phosphorylation of SYK and BTK (**Figures 5J, 5K, S5D**). Similar findings were observed in I29 cells (**Figures 3L, 3M, S5E**).

Both IgA+ B cell lines were characterized by higher basal phosphorylation level (**Figures 5I, S5F, S5G**) which was dependent on the BCR signaling, as treatment with the Syk inhibitor, R406, reduced phosphorylated proteins including pBTK while leaving pSYK unchanged (**Figures 5N, 5O, S5H-J**). To determine whether IgA BCR mediates stronger BCR signaling in human B cells as well, we stimulated sorted GC from tonsils and observed that IgA+ GC B cells had increased total phosphorylated proteins compared to IgM+ GC cells (**Figures 5P, 5Q**). Our findings highlight that IgA BCR mediates stronger antigen receptor signaling in both basal and activated states, a property also conserved in human.

### IgA resistance to FasL-dependent cell death increase GC competitiveness

In the LZ compartment, in addition to providing help and positively selecting B cells, T cells can eliminate B cells via activation of the FasL-Fas-dependent cell death pathway.^34^ Previous reports suggested that different B cell isotypes might be differently sensitive to Fas-mediated killing^63,64^, but the role of IgA BCR in this process has not been investigated. Therefore, we sought to test whether IgA BCR confers resistance to Fas-mediated death using CH12 cells, which express similar levels of Fas regardless of isotype (**Figure S6A**) and FasL expressing vesicles, which express the physiological FasL trimer.^65^ IgA+ CH12 were significantly protected from Fas dependent death compared to IgM+ CH12, suggesting an IgA intrinsic basal protection, likely driven by tonic BCR signaling (**Figures 6A, S5C**).

**Figure 6.**
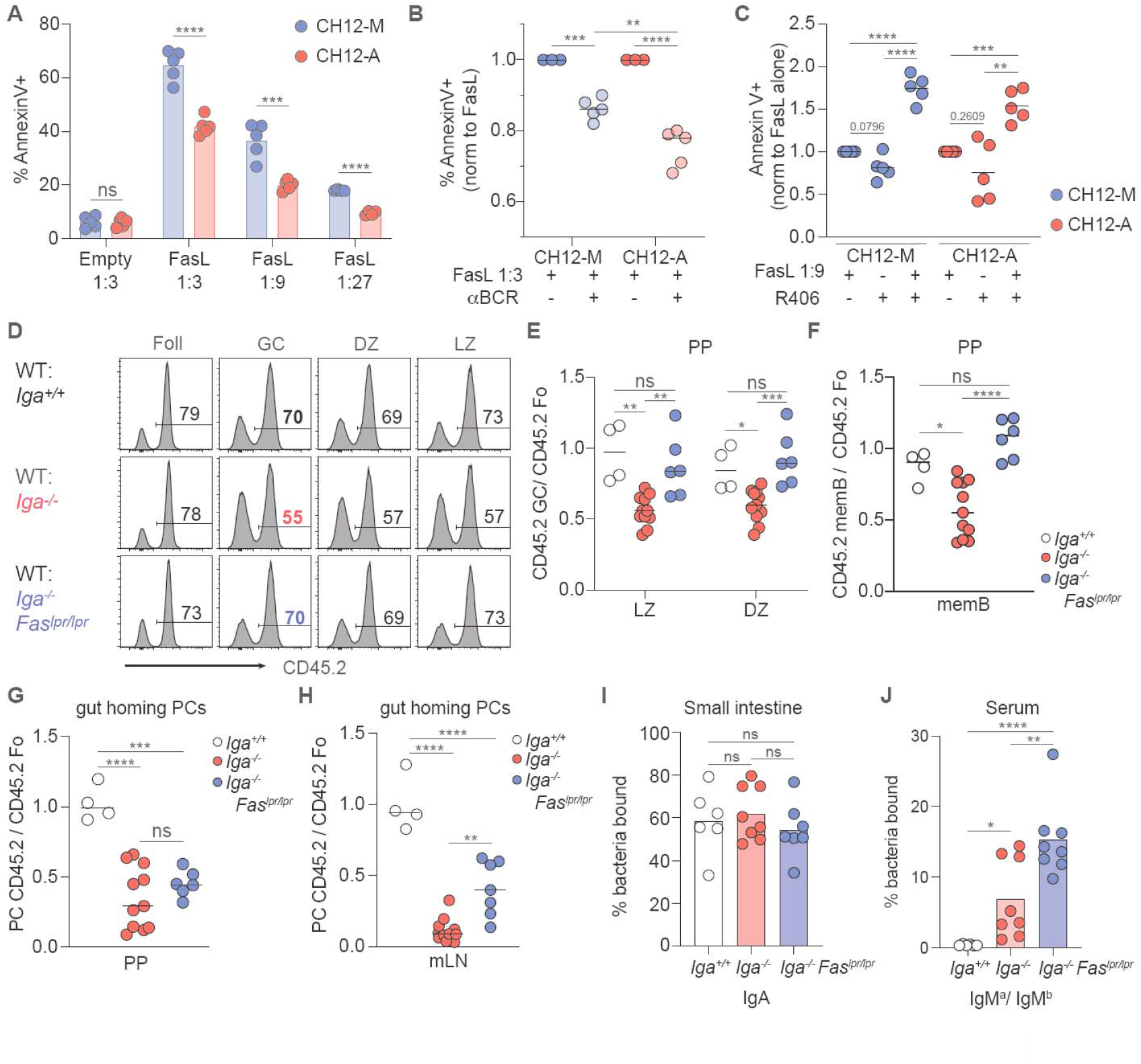
IgA+ GC B cells are more resistant to Fas-dependent counter selection. (A) Percentage of dead (defined as annexin V+) CH12 cells 5 hours after incubation with FasL vesicles at indicated dilutions. Representative of at least 3 independent experiments. (B) Percentage of dead CH12 cells after BCR stimulation and incubation with FasL vesicles. Data is normalized to FasL vesicles alone. Compiled data from at least 3 independent experiments. (C) Percentage of dead cells after incubation with R409 and FasL vesicles. Data is normalized to FasL vesicles alone. Compiled data from 3 independent experiments. (D) Representative histograms of CD45 staining on mBM chimeras for indicated B cell populations. (E) Ratio of frequency of CD45.2 LZ or DZ B cells to CD45.2 Foll B cells in PP of mBM chimeras. (F) Ratio of frequency of CD45.2 memory B cells to CD45.1 memory B cells from PPs of mBM chimeras. (G, H) Ratio of frequency of CD45.2 gut homing PCs to CD45.1 gut homing PCs from PPs(G) and mLN(H) of mBM chimeras. E-H are compiled data from 3 independent experiments with 1-3 mice per group. (I, J) mBM chimeras were made in uMT hosts to track antibodies using allotypic isotypes. (I) Percentage of endogenously coated small intestinal bacteria with IgA antibodies. (J) Percentage of intestinal commensal from uMT mice stained by IgM^a^ or IgM^b^ antibodies from mBM serum. Percentage shown as IgM^a^/IgM^b^ ratio. I, J are compiled data from 3 independent experiments with 2-3 mice per group. Unpaired t-test was used in a. One-way anova was used in b, c, e-j. ns=not significant, *p<0.005, **p<0.001,***p<0.0005, ****p<0.0001.

To test how BCR signaling may offset Fas signaling, CH12 B cells were stimulated immediately prior to incubation with FasL vesicles. While BCR stimulation rescued both CH12-M and CH12-A from Fas-mediated apoptosis, a significantly higher degree of protection was achieved in IgA+ CH12 (**Figure 6B**). Conversely, inhibition of BCR signaling with R406 enhanced CH12-M and CH12-A sensitivity to FasL vesicles, with CH12-A showing increased propensity to undergo cell death (**Figure 4C**). Together this data suggests that IgA BCR signaling provides protection against Fas mediated death initiated by membrane FasL, suggesting an intrinsic mechanism that might underpin IgA GC B cell survival in vivo.

Cell-intrinsic suppression of GC B cell survival via FasL-Fas has been shown to be important in preventing outgrowth of GC; inability to undergo FasL-mediated cell death predisposes the host to autoimmunity and B cell lymphoma.^34,66–68^ Interestingly, such constraints on GC B cell accumulation in vivo is tissue specific: while Fas-dependent deletion is a dominant mechanism to maintain GC in peripheral lymph node and mLN, in PPs, GC B cells resistance to Fas killing did not confer a competitive advantage.^34^ In PPs, most GC B cells carry an IgA BCR, thus IgA+ GC B cells might be equipped with molecular machinery that permits a higher threshold for Fas-dependent cell death. Since GC B cells express similar levels of surface Fas independently from the isotype in PPs, (**Figure S6B**), we speculated that *Iga*^*-/-*^ B cells are preferentially killed via Fas and therefore could be rescued by inactivation of the Fas-induced cell death pathway. To test this hypothesis, we generated mixed BM chimeras with BM from WT and *Iga*^*-/-*^, or *Iga*^*-/-*^ resistant to Fas-mediated cell death (*Iga*^*-/-*^ Fas^LPR/LPR^). In PPs, we observed that, in contrast to *Iga*^*-/-*^ B cells, *Iga*^*-/-*^ Fas^LPR/LPR^ B cells were able to compete with WT B cells during GC reaction in both LZ and DZ (**Figures 6D, 6E**), and their restored GC fitness led to physiologic memory B cell compartment formation (**Figure 6F**). In mLN, where a very small number of GC B cells express IgA BCR, resistance to Fas-mediated cell death allowed *Iga*^*-/-*^ Fas^LPR/LPR^ B cells outgrowth as GC B cells and memory B cells, supporting previous findings^34^ (**Figures S6C, S6D**). No preferential isotype expansion was observed in *Iga*^*-/-*^ Fas^LPR/LPR^ B cell subsets (**Figures S6E, S6F**), suggesting that the GC competitive ability was not driven by B cells carrying a specific isotype, but was instead the result of global rescue of GC B cells from Fas-mediated cell death. Interestingly, although *Iga*^*-/-*^ Fas^LPR/LPR^ GC B cells could generate PCs at a rate similar to WT B cells (**Figure S6G**) they were nevertheless unable to differentiate into gut homing PCs (**Figures 6G, 6H**) highlighting a previously undescribed role for the IgA BCR in mucosal imprinting that is uncoupled from GC selection. In line with this finding, mixed BM chimeras generated in μMT hosts using allotypic donors (IgHa vs IgHb) to track Ab origins revealed that endogenous intestinal bacteria were coated with IgA coming from WT B cells (**Figure 6I**), with negligible contribution of IgM from *Iga*^*-/-*^ or *Iga*^*-/-*^ Fas^LPR/LPR^ (**Figure S6H**). In contrast, *Iga*^*-/-*^ Fas^LPR/LPR^ serum gave rise to significantly more IgM specific to commensals (**Figure 6J**), likely due to B cell outgrowth in mLN (**Figures S6C, S6F**). These results indicate that Fas counterselection is critical for IgA+ GC B cells’ enhanced competition and point to an IgA-intrinsic resistance to Fas-dependent cell death in vivo. However, rescue of non-IgA GC B cells by Fas deletion does not increase the generation of gut-homing PCs, suggesting that additional mechanisms in IgA GC B cells imprint the ability to direct PCs to the intestinal mucosa.

## Discussion

Class switch recombination to IgA, and the consequent acquisition of surface IgA, has been mainly viewed as a default process for B cells, enforced by the abundant active TGFβ available in the sub-epithelial dome of PPs to recently activated B cells.^26,27^ This process imprints the B cell response to intestinal antigen for the generation of mostly IgA-secreting plasma cells. The incorporation of J-chain for rapid retrotranslocation via pIgR^69,70^, the dimeric form for increased avidity^71^ and the somatic mutation loads^16–20^ render IgA the most efficient mucosal antibody isotype. Thus, induction of the IgA response was viewed as a process that generates essential antibodies for secretion in the intestinal lumen most efficiently.

Here we described an additional role of IgA during the GC B cell reaction, providing a new perspective on the evolutionary requirements of IgA class switch in B cells. Together, IgA is critical to promote fitness during the GC reaction and imprint gut-homing properties on nascent plasma cells. The conflicting demands of broad IgA reactivity and affinity-based GC selection has been partially reconciled by recent works describing tunable affinity maturation in Peyer’s patches, but the mechanism underpinning the GC reaction to intestinal antigens have not been elucidated. Interestingly, lower microbiome complexity that restores affinity maturation efficiency and leads to enhanced clonal selection in Peyer’s patches germinal center is associated with a loss of IgA+ GC B cells^36^, suggesting IgA BCR tunes GC selection toward a more permissive reactivity. How, then, are IgA+ B cells selected during GC reaction to generate a humoral response at the intestinal barrier that can simultaneously coat several classes of commensals? We propose a model in which the IgA BCR is inherently required for the generation of the broad mucosal antibody repertoire. IgA+ GC B cells display intrinsic protection from apoptosis that is centered around an enhanced BCR intracellular signaling and increased T cell-derived positive cues during selection of antigen-specific clones. Mechanistically, IgA BCR allows GC B cells to offset the Fas-FasL cell death signaling, therefore allowing low affinity B cells to persist and contribute to the plasma cell compartment.

The structural basis for the superior positive selection of IgA GC B cells remains unclear. While reports exist that the intracellular tail of IgG1 and IgE do not contribute to their intrinsic effect on GC positive selection, the IgA intracellular tail is distinct from all the other isotypes both in terms of length and sequence and could underpin the specific effect on IgA+ GC B cells during responses in PPs. Additionally, it also conceivable that Ig isotype-specific membrane and constant regions could play a role in IgA expressing cells, by either facilitating BCR clustering and thus lowering the threshold of B cell selection in the GC, or by altering BCR stiffness and thus rendering IgA+ GC B cells superior in extracting antigen displayed on FDCs.^72,73^

In our model, strong IgA BCR signaling offsets FasL-mediated cell death signals and promotes GC competition. In contrast to IgA, IgE BCR mediates autonomous, sustained calcium signaling which constrains GC formation and instead promotes plasma cell differentiation.^42,45^ Deletion of Fas increases the ability of IgA-deficient cells to compete in the GC reaction, while Fas-deficient B cells generate a high number of IgE+ PC.^68^ One possible explanation for these apparently contradicting results is that BCR signaling is integrated to a different extent by different isotype carrying GC B cells: IgE having the strongest signaling does promote PC formation in line with a strength of signaling model^31,74^, while IgA BCR signaling, while dominant over the other BCR isotypes present in PP GC, allows for both GC competition and PC differentiation. Thus, the high IgE titer observed in the Fas or FasL deficient mice might be due to expansion of IgG1+ GC B cells that sequentially switch to IgE and then contribute to the humoral immunity. Moreover, since IgE B cells do not efficiently participate in the GC reaction, it is unlikely that they are sensitive to Fas/FasL level at that stage. The stricter FasL-mediated counterselection of low affinity non-IgA GC B cells is an appropriate strategy to avoid the generation of rogue, autoreactive B cell clones.^34,68^

We also discovered that the IgA BCR is required in a competitive setting for the efficient upregulation of migratory receptors CCR9 and α4β7 and eventual homing of PCs to the lamina propria. Surprisingly, this process is independent of Fas-FasL, as restoring competitiveness of non-IgA+ B cells during GC reaction had no effect on their ability to upregulate CCR9 and α4β7. Thus, additional step(s) that control functional generation of intestinal humoral immunity must take place in PPs during the response to intestinal antigen. While the homing ability could be imprinted during LZ selection in the GC, it could also be imprinted on plasmablasts and even fully differentiated PCs, which can maintain BCR surface receptor expression^75,76^, suggesting that BCR signaling could shape PC homing outside of the GC reaction.

Interestingly, IgA plasma cells isolated from lungs and mediastinal lymph nodes of mice infected with influenza also show marked upregulation of mucosal homing receptors^77^, suggesting a direct link between IgA BCR and PC migratory capabilities that is independent of the niche that generates PCs. Which signals, and at which step of PC differentiation, control the upregulation of mucosal homing receptors in PCs remain to be elucidated. We described the existence of a process that equips IgA+ PCs with gut homing receptors during responses to commensals in order to direct antibody production into the intestinal lamina propria; however, whether such process remains in place or are altered during enteric infection or dysbiosis, situations that can lead to non-IgA+ PC homing to the lamina propria,^78–80^ will require additional studies.

Given the dynamic impact of IgA on gut autoinflammation, understanding how IgA B cells are selected in the GC and give rise to IgA intestinal responses could be exploited therapeutically to develop strategies that restore commensal homeostasis. Moreover, elucidating how surface BCR signaling during the GC reaction regulates plasma cell tissue specification could be used to direct isotype-specific secreting plasma cells to different barrier tissues, including the upper respiratory and genital tract, and to generate vaccines against mucosal pathogens.

