## Supplement for "Antigen receptor signaling and cell death resistance controls intestinal humoral response zonation"

**ACKNOWLEDGMENTS**
We thank Carol Schrader for the I29μ cell line, Stephanie Moses for FasL vesicle preparation, Garnett Kelsoe for generously allowing use of NB21 cells, and Kelsey Howley and Carly Burke for mouse genotyping and colony maintenance. We thank E. V. Dang for critical reading of the manuscript.
This work was supported by National Institutes of Health (NIH) grant no. AI155727 and no. R21AI173903 (to A.R.) and NIH training grants T32 AI132152 and T32 AI007349 (to F.R.). J.R.M. is supported by the Intramural Research Program of the National Institutes of Health, Center for Cancer Research, National Cancer Institute. We acknowledge the University of Massachusetts Flow Cytometry Core Facility for access to sorting services.

AUTHOR CONTRIBUTIONS
F.R. and A.R. conceived the study, developed the concept, designed the experiments, and wrote the paper. F.R. performed the experiments, analyzed the data, and interpreted the results. S.L., M.J.W. and O.B. performed experiments and analyzed the data. G.B. provided *Iga*$^{-/-}$ mice. All authors contributed to the review and editing of the manuscript. A.R. reviewed the data and supervised the research.

### MATERIALS AND METHODS

#### Mice

C57BL/6J (CD45.2) (Stock No: 000664), Ly5.2 (CD45.1) congenic B6.SJL-Ptprca Pepcb/BoyJ (Stock No 002014), Rosa26-flox-stop-flox-tdTomato (Stock No: 007914), Aicda-cre (Stock No: 007770), B6.MRL-Fas$^{lpr}$ (Stock No: 000482), B6J.Cg-Gt(ROSA)26Sor$^{tm95.1(CAG-GCaMPgf)Hze}$/MwarJ (gCAMP) (Stock No:028865), B6.Cg-Tg(TcraTcrb)425Cbn/J (OTII, Stock No:004194), B6.Cg-Rag2$^{tm1.1Cgn}$/J (Stock No: 008449), B6.129SC-*Ighm*$^{tm1Cgn}$/J were purchased from the Jackson Laboratory. Gregory Barton provided *Iga*$^{-/-}$ mice, IgMa mice were from an internal colony. India mice were from NIH (Mayer et al. Science 2017). cMYC-GFP mice were from Oliver Bannard. All mice at UMass Chan Medical School were bred and maintained under standard 12:12 hours light/dark conditions and housed in specific pathogen-free (SPF) conditions. Female and male mice were analyzed at 8-12 weeks of age and littermates of the same sex were cohoused and randomly assigned to groups. All procedures were conformed to ethical principles and guidelines approved by the UMass Chan Medical School Institutional Animal Care and Use Committee.

#### Bone marrow chimeras, retroviral transduction

Ly5.1/2, WT, or uMT mice were lethally irradiated twice with 550 rads gamma-irradiation, 3 hours apart, then intravenously injected with 1-3 x $10^6$ bone marrow cells. Bone marrow was harvested by flushing both tibia and femurs of donor mice, counted, and mixed at ratio indicated in text. Mice were analyzed 8-12 weeks after cell transfer.

For retroviral transduction, PlatE cells were transfected with murine stem cell virus retroviral constructs encoding full length Bcl2 with Lipofectamine 2000 (Invitrogen) following manufacturer's protocol. For transduction of BM-derived cells, BM was harvested 4 days after 5-flurouracil (Sigma) injection and cultured in the presence of recombinant IL-3, IL-6, and mouse stem cell factor (100 ng/ml, Peprotech). BM cells were spin-infected twice with a retroviral construct expressing Bcl2 with GFP as a reporter. One day after the last spin infection, the cells were injected into lethally irradiated Ly5.1/2 recipients. PPs and mLN were analyzed 8-10 weeks later.

#### CT immunization and Salmonella infection

For cholera toxin responses, mice were immunized with 10 ug of cholera toxin (Sigma) in PBS by oral gavage. Animals received cholera toxin every 7 days for three consecutive weeks. Memory B cells were sorted and analyzed 7 days after last immunization.

The SL1344 metabolically defective (*aroA*-) S. typhimurium strains were provided by Milena Bogunovic (UMass Chan) and grown at 37C in Luria broth supplemented with appropriate antibiotics to preserve mutation and plasmid. To induce Ova expression, IPTG was added at an $OD_{600}$ of 0.4 and allowed to grow for another 2 hours. For the Salmonella infection, wildtype mice were orally gavage three times, on alternate days with $10^9$ CFUs of *aroA*- Salmonella expressing Ova in 200 ul 5% sodium bicarbonate. For antigen presentation experiments, three days after the last dose, follicular and germinal center B cells were sorted for from PPs. For mixed BM chimera infections, 8 weeks after BM reconstitution, mice were infected as described above. Seven days after the third dose of Salmonella, mice were scarified and B cell populations were assessed by flow cytometry.

**Sample Preparation and Flow Cytometry**

Peyer's patches and mesenteric lymph nodes were collected into complete media (RPMI, 5% heat inactivated FBS, 10 mM HEPES, 1%Pen/Strep). The tissue was then smashed through a 70 micron cell strainer. Isolated cells were then washed with FACS buffer (1X DPBS, 2% heat inactivated FBS, 2mM EDTA). Cell suspensions were stained with LIVE/DEAD Fixable Aqua Dead Cell Stain (Invitrogen) or Fixable Viability Dye eFluor780 (Invitrogen) in FACS buffer, and Fc receptors blocked with anti-mouse CD16/32 (2.4G2). BD Cytofix/Cytoperm kit (BD BDB554714) was used for fixation and intracellular staining. Cells were incubated for 20 min on ice with antibodies described in table S1. Data were collected on a BD SORP or LSR II and analyzed in FlowJo v10.7 software.

For cell sorting, cells were stained as described above and sorted on a BD FACSAria II with an 85-micron nozzle. Cells were maintained at 4C until sorting. Sorted PP Follicular and germinal center B cells were used for antigen presentation. PP memory B cells were used for NB21 co-culture. Germinal center B cells were sorted as live IgD- GL7+ CD38- CD138- isotype+. Follicular and memory B cells were sorted as live IgD+ GL7- CD138- or live IgD- GL7+ CD38+ CD138- CD73+ CD80+ respectively into complete media for further culture.

For BrdU incorporation experiments, animals were given 2.5 mg of BrdU in a single i.p. injection and sacrificed 30 minutes or 3.5 hours later. Staining was performed using the FITC BrdU Flow Kit (BD PharMingen) according to the manufacturer's instructions. To stain for intracellular antigens, cells were first stained for surface markers and then fixed and permeabilized using the BD Cytofix/Cytoperm Kit per the manufacturer's instructions.

For anti–active caspase-3 staining, cells were maintained on ice during harvesting and processing. Cells were stained for with biotin–conjugated anti–active caspase-3 (BD Biosciences) following fixation and permeabilization according to the manufacturer's instructions and subsequently stained with FITC or AF647 streptavidin.

Bacterial flow cytometry was adapted from Koch et al 2016 Cell with minor adjustments described here. Bacteria were washed in sterile-filtered DPBS with 1% bovine serum albumin (BSA, Fisher) and resuspended at approximately $5 \times 10^6$ bacteria/mL. Mouse serum was diluted 1:25 in PBS/BSA buffer and 25uL of this solution was mixed with 25uL diluted microbes in u-bottom plate. Staining was performed with purified or fluorochrome conjugated antibodies. Cells were washed and resuspended in 1% BSA in DPBS with SYBR Green (Invitrogen) and analyzed by FACS.

**Cell culture**

The I29μ cell line was obtained from Carol Schrader (UMass Chan). The CH12 cell line was obtained from Jagan Muppidi (NIH). NB21 cells were a gift from G. Kelose (Duke).

Both CH12 and I29 were cultured in B cell media (10% FCS, 100 units/mL penicillin, 100 units/mL streptomycin, MEM nonessential amino acids, 10mM HEPES, 1mM sodium pyruvate, 55mM 2-Mercatoethanol). To induce class switch to IgA, CH12 and I29 cells were cultured for 48 hours with 2ng/ml TGFβ and 50 ug/ml LPS. The IgA+ population was sorted with a BD FACS Aria II and maintained as CH12 IgA and I29 IgA cell lines.

Freshly isolated memory B cells from Peyer's patches were FACS sorted as described above and cocultured with NB21 feeder cells as previously described with adjustments (Kuraoka et al., 2016; Stewart et al., 2018). Briefly $15 \times 10^3$ NB21.2D9 cells/ well were  seeded into 96-well u bottom plates in 100 ul of B cell media and grown at 37C and 5% $CO_2$ the morning of B cell sorting and coculturing. 2000 memory B cells in 100 ul B cell media were added to each well. Culture plates were centrifuged at day 3.5 and the supernatant collected for Ig detection.

**FasL killing Assay**

FasL vesicles were prepared as previously described (Jodo et al 2000 J Immunol). Briefly, FasL vesicles were collected from N2-mFasL cells that overexpress mouse FasL lacking the metalloproteinase-sensitive site and empty vesicles from Neo-N2 cells that did not contain FasL used as control. When cells reached ~70% confluence, fresh culture medium without G418 replaced the G418-containing medium. 48 hours later, the supernatants were collected and centrifuged at 4°C, 13,000 rpm in a Sorvall Superspeed centrifuge for 30 min to remove cell debris. Next, the cell-free supernatants were centrifuged for 3 hours at 4°C at 25,000 rpm in a Beckman ultracentrifuge with a SW25 rotor. Following ultracentrifugation, the murine FasL vesicle pellet was suspended with culture medium to 7% of the original volume and passed through a 0.45-mm sterile filter and used for cytotoxicity assays.

For the in vitro FasL killing, 200,000 CH12 cells were plated in 50 uL of warmed calcium media in u-bottom 96 well plates. FasL or empty vesicles were diluted in warmed calcium media at the dilutions indicated in the figures and 50ul were added to each well. Cells were incubated at 37C with 5% CO2 for 5 hours. Cell death was assessed with annexin V and DAPI. For annexin V staining, cells were stained according to the manufacturer's instructions (BD cat 556419). DAPI was added after annexin V staining right before cells were run at the FACS.

**B cell Stimulation**

For calcium stimulation, Peyer's patches were harvested into warmed calcium media (DMEM with 2.5% FCS and 10mM HEPES) and processed as described above. Germinal center and memory B cells were enriched for using EasySept negative B cell enrichment with modifications. Briefly, PPs were incubated with biotinylated antibodies against IgD, CD3, CD4, CD8, and CD138. Following negative enrichment, experiments using calcium dyes were incubated with surface antibodies GL7 and CD38 and 6uM FuraRed (Invitrogen F-3020) and 3uM Fluo-3 AM (Invitrogen F23915) at 37C for 20 minutes. Cells were washed and resuspended in warmed calcium media and rested 15-30 minutes at 37C prior to stimulation. For gCAMP experiments, after negative B cell enrichment, cells were stained with GL7 and CD38 at room temperature for 10 minutes, washed and rested at 37C for 10-30 minutes before stimulation. In some experiments cells were incubated with 100 nM Ibrutinib for 10 minutes at room temperature. For all experiments done with gCAMP mice or calcium dyes, cells were run on the FACS for 30 seconds to collect baseline and then stimulated with 10 ug/mL anti-BCR. After 2-3 minutes 1ug/mL Ionomycin was added to measure maximum calcium flux.

For CH12 and I29 BCR stimulation, 200,000 B cells were resuspended in 400 ul of warmed calcium media and rested for 30 minutes prior to stimulation. Cells were stimulated with 10ug/mL anti-BCR in warm calcium media for 1 minute then fixed for western blot or phospho-flow. For western blots, cells were spun at max speed RT for 10 seconds, the supernatant was removed, and the cell pellet was lysed in RIPA buffer (Thermo Cat 89900) with phosphatase inhibitor (Thermo Cat 1861281). Lysed cells were left on ice for 30 minutes then spun down at max speed for 10 minutes. The supernatant was used as whole cell lysates for western blots. Whole cell lysates were boiled for 5 minutes in Laemmli sample buffer and 2-ME then run on 10% tris-glycine cells. Proteins were transferred to PVDF membrane and blocked with 2% BSA in TBST for one hour at RT then incubated with anti-mouse phosphotyrosine-HRP antibody (Cell Signaling) overnight at 4C. Blots were washed with TBST followed by development with SuperSignal West Pico PLUS chemiluminescent Substate (Thermo Cat 34577). Actin was detected with mouse anti-actin (Cell Signaling) for 1.5 hours at room temperature followed by anti-mouse HRP (Biolegend) for 1.5 hours and detected with ECL Prime Western blotting detection reagent (Cytiva Cat RPN2232). ImageJ software was used for band densitometry.

For phosphoflow, after anti-BCR stimulation, cells were fixed with 16% PFA at a final concentration of 1% PFA for 10 minutes at room temperature. Cells were pelleted and resuspended in 1 mL of ice-cold methanol and left overnight at -20C. The following day, cells were washed with 3 ml of FACS buffer and then 4 ml of FACS buffer. Cells were blocked in a u-bottom 96 well plate for 20 minutes with Fc block at room temperature, then stained with phosphor-tyrosine (Biolegend) or phosphoBTK (Biolegend) for 45 minutes at room temperature and then analyzed by BD SORP or LSR II.

For Syk inhibition, 200,000 I29 or CH12 cells were incubated at 37C in calcium media containing 2uM R406 Syk inhibitor for 45 minutes. Cells were washed with warm calcium media and pelleted. Cells were resuspended in warm calcium media and immediately fixed for phosphoflow as described above.

For antigen presentation assays, follicular, IgM+, IgG+ and IgA+ germinal center B cells were sorted from Peyer's patches as described above. OT-II T cells were isolated from spleens of OT-II Rag KO mice. B cells and T cells were co-cultured 1:1 in 200 uL of B cell media in 96 well u-bottom tissue culture plate and incubated at 37C for 18 hours. Follicular B cells were cultured with or without 2 uM Ova protein. CD69 expression on T cells was assessed by flow cytometry.

**Immunohistochemistry and Immunofluorescence**

For immunohistochemistry, acetone fixed cryosections were stained with the following antibodies: goat purified anti-mouse IgD and biotin anti mouse GL7. Then sections were stained with the secondary antibodies anti-goat HRP and streptavidin Alkaline phosphatase (AP).

For immunofluorescence, tissues were fixed in 4% paraformaldehyde in PB overnight at 4°C, washed three times for 30 min in PB, then moved to 30% sucrose in PB overnight. Tissues were flash frozen in Tissue-Tek Cryomold (VWR) the next day, and 7-mm sections were cut and then dried for at least 1 hour before staining. Sections were rehydrated in PBS with 1% bovine serum albumin (BSA) for 10 min and then stained overnight at 4C and stained for subsequent steps at room temperature for two hours, all in PBS with 1% BSA, 2% mouse serum, 2% rat serum and 2% donkey serum. Sections were stained with primary antibodies: Goat anti-mouse IgD (goat polyclonal GAM/IgD(FC/7S, Cedarlane Labs), rabbit anti-mouse RFP and Alexa647-conjugated anti-CD35/21. Sections were then stained with the secondary antibodies: Cy3 anti-rabbit, AF488 anti-goat, and 40,6-diamidino-2-phenylindole (DAPI). tdTomato cells were counted in the LZ (CD35+) and DZ (CD35-) regions.

**Human Tonsils**

Human tonsil cells were thawed in 37°C IMDM media with 10% Fetal Bovine Serum (FBS), 1% Glutamax, 1% HEPES, 1% Penicillin/Streptomycin and 0.1% β-mercaptoethanol. Single cell suspensions were blocked with Fc Block at 4°C for 10min and subsequently, stained with CD19 PE-Cy7, CD27 BV605, CD10 APC or APC-Cy7, and CD138 PerCP-Cy5.5 antibodies to identify Germinal Center B cell and Follicular B cells. For sorting of IgA+ GC B cells without stimulating the desired population, cells were incubated IgG FITC and IgM PE, and the double negative population was sorted with FACS Aria (BD Bioscience). For sorting of IgM+ GC B cells without stimulating the desired population, cells were labeled with additional IgG FITC and IgA PE and sorted as above.

For the western blot signaling assay, sorted cells were rested and stimulated with 10ug/ml anti-IgA or anti-IgM for 5 min. Cells were lysed for 30 min on ice in RIPA buffer containing 0.5% dithiothreitol (DTT) and protease inhibitor cocktail. Lysates were centrifuged for 10 min at 4°C at 14,000g and supernatant was collected. Proteins were separated by electrophoresis using NuPage-Bis-Tris gels (Invitrogen) and blotted onto polyvinylidene fluoride membranes. Non-specific binding was blocked with 5% BSA in TBS-Tween (0.1%) followed by incubation with anti-phosphotyrosine overnight at 4 °C and secondary antibody horseradish peroxidase (HRP) conjugated antibodies for 1 hour at room temperature. Membranes were washed thoroughly with TBS-Tween (0.1%) after antibody incubations and developed using ECL reagents (Millipore). For re-probing, blots were stripped for 10 min at RT with Restore PLUS stripping buffer (Thermo Fisher Scientific). Blots were then incubated with β-actin primary antibody and secondary antibody HRP-conjugated antibody for 1 hr at RT, washed with TBS-Tween, and developed using ECL reagents (Cytiva Amersham). ImageJ software was used for band densitometry.

**Statistical analysis and reproducibility**

Statistical analyses were performed using GraphPad Prism v8.0 using two-way ANOVA with Bonferroni's multiple comparisons test or unpaired t test. Data are presented as means ± SEM except Figure 3B and 3C are SD. Differences between group means were considered significant at indicated p value. The significance values are stated in all graphs and the number of biological replicates (n) is stated in the figure legends.

Table S1. Flow Cytometry antibodies

| Antibody | Dilution | CAT | Vendor | RRID |
|---|---|---|---|---|
| Purified Fc Block (CD16/32) | 1:200 | 101330 | Biolegend | AB_2561482 |
| B220 BV711 | 1:1000 | 103255 | Biolegend | AB_2563491 |
| CD19 AF700 | 1:1000 | 115528 | Biolegend | AB_493735 |
| CD45.1 AF700 | 1:100 | 110724 | Biolegend | AB_493733 |
| CD45.1 Pacific Blue | 1:100 | 110722 | Biolegend | AB_492866 |
| CD45.1 BV650 | 1:100 | 110736 | Biolegend | AB_2562564 |
| CD45.2 AF700 | 1:100 | 109822 | Biolegend | AB_493731 |

| | | | | |
|---|---|---|---|---|
| CD45.2 Pacific Blue | 1:100 | 109819 | Biolegend | AB_492873 |
| CD45.2 BV650 | 1:100 | 109836 | Biolegend | AB_2563065 |
| IgD BV510 | 1:1000 | 405723 | Biolegend | AB_2562742 |
| IgD PerCP Cy5.5 | 1:1000 | 405710 | Biolegend | AB_1575113 |
| IgD Pacific Blue | 1:1000 | 405712 | Biolegend | AB_1937244 |
| IgD Alexa Fluor 647 | 1:1000 | 405708 | Biolegend | AB_893528 |
| CD138 BV421 | 1:1000 | 142508 | Biolegend | AB_11203544 |
| GL7 AF647 | 1:1000 | 144606 | Biolegend | AB_2562185 |
| GL7 PerCP Cy5.5 | 1:500 | 144610 | Biolegend | AB_2562979 |
| CD38 PE Cy7 | 1:1000 | 102718 | Biolegend | AB_2275531 |
| CD73 PerCP Cy5.5 | 1:100 | 127214 | Biolegend | AB_11219403 |
| CD80 BV605 | 1:100 | 104729 | Biolegend | AB_11126141 |
| CD86 BV605 | 1:100 | 105037 | Biolegend | AB_11204429 |
| CD86 FITC | 1:100 | 105005 | Biolegend | AB_313148 |
| CXCR4 Biotin | 1:100 | 551968 | BD Biosciences | AB_394307 |
| CXCR4 PE | 1:100 | 551966 | BD Biosciences | AB_394305 |
| CCR9 PE | 1:100 | 129708 | Biolegend | AB_2073249 |
| LPAM-1 biotin | 1:100 | 120612 | Biolegend | AB_11203892 |
| Streptavidin AF488 | | 405235 | Biolegend | |
| Streptavidin PerCPCy5.5 | | 405214 | Biolegend | |
| Streptavidin BV605 | | 405229 | Biolegend | |
| Streptavidin BV650 | | 405231 | Biolegend | |
| Streptavidin PE Cy7 | | 25-4317-82 | eBioscience | |
| IgM APC | 1:100 | 550676 | BD Biosciences | AB_398464 |
| IgM FITC | 1:100 | 553437 | BD Biosciences | AB_394857 |

| | | | | |
|---|---|---|---|---|
| IgM Biotin | 1:100 | 553436 | BD Biosciences | AB_394856 |
| IgMa FITC | 1:100 | 408606 | Biolegend | AB_940541 |
| IgMa PE | 1:100 | 408608 | Biolegend | AB_940545 |
| IgMb FITC | 1:100 | 406206 | Biolegend | AB_315039 |
| IgMb PE | 1:100 | 406208 | Biolegend | AB_315041 |
| IgG1 FITC | 1:1000 | 406605 | Biolegend | AB_493292 |
| IgG1 PE | 1:1000 | 406607 | Biolegend | AB_10551439 |
| IgG2b FITC | 1:1000 | 406705 | Biolegend | AB_493296 |
| IgG2b PE | 1:1000 | 406708 | Biolegend | AB_2563381 |
| IgG3 biotin | 1:100 | 406803 | Biolegend | AB_315070 |
| IgA FITC | 1:1000 | 1040-02 | Southern Biotech | AB_2794370 |
| IgA PE | 1:1000 | 1040-09 | Southern Biotech | AB_2794375 |
| IgA biotin | 1:1000 | 1040-08 | Southern Biotech | AB_2794374 |
| Rabbit anti-active Caspase 3 biotin | 20ul per test | 550557 | BD Biosciences | AB_393750 |
| Kappa AF700 | 1:500 | 409508 | Biolegend | AB_2563583 |
| CD22 APC (Clone OX-97) | 1:500 | 126110 | Biolegend | AB_2561630 |
| I-A/I-E PerCP Cy5.5 | 1:500 | 107626 | Biolegend | AB_2191071 |
| CD79b PerCP CY5.5 | 1:500 | 132810 | Biolegend | AB_2632918 |
| CD69 PE Cy7 | 1:500 | 104512 | Biolegend | AB_493564 |
| CD4 Alexa Fluor 700 | 1:500 | 116022 | Biolegend | AB_2715958 |
| Vα2 PE | 1:200 | 553289 | BD Biosciences | AB_394760 |
| TCRβ Pacific Blue | 1:500 | 109226 | Biolegend | AB_1027649 |
| Fas PE-Cy7 | 1:500 | 557653 | BD Biosciences | AB_396768 |

| | | | | |
|---|---|---|---|---|
| Annexin V FITC | 5 ul per test | 556419 | BD Biosciences | AB_2665412 |
| Phospho-BTK (pY223) PE | 1:100 | 562753 | BD Biosciences | AB_2737769 |
| Phospho-Syk (pY352) AF647 | 1:100 | 557817 | BD Biosciences | AB_396884 |
| phospho-tyrosine PE | 1:100 | 309310 | Biolegend | AB_2572196 |
| Live/Dead eFluor780 | 1:2000 | 4302687 | BD Biosciences | |
| Live/Dead eF450 | 1:500 | 65-0863-18 | eBioscience | |
| Human Fc Block | | 422302 | BD Biosciences | |
| Anti-human CD19 PE Cy7 | 1:100 | 560728 | BD Biosciences | AB_1727438 |
| Anti-human CD27 BV605 | 1:67 | 302830 | BD Biosciences | |
| Anti-human CD10 APC | 1:100 | 312210 | BD Biosciences | |
| Anti-human CD10 APC-Cy7 | 1:100 | 312212 | BD Biosciences | |
| Anti-human CD138 PerCP-Cy5.5 | 1:100 | 356510 | BD Biosciences | |
| Goat F(ab')2 Anti-Human IgM-PE | 1:200 | 2022-09 | Southern Biotech | AB_2795614 |
| Goat F(ab')2 Anti-Human IgA-PE | 1:200 | 2052-09 | Southern Biotech | AB_2687523 |
| Goat F(ab')2 Anti-Human IgM-unlabeled | | 2022-01 | Southern Biotech | AB_2795610 |
| Goat F(ab')2 Anti-Human IgA-unlabeled | | 2052-01 | Southern Biotech | AB_2795709 |
| Goat F(ab')2 Anti-Human IgG-FITC | 1:200 | AHI1308 | Thermo Fisher | AB_1500723 |

Other Reagents

| | | | | |
|---|---|---|---|---|
| Stop Solution for TMB Substrate | | 423001 | Biolegend | |
| IgD Biotin | 1:1000 | 405734 | Biolegend | **AB_2563344** |
| stop solution | | 77316 | Biolegend | |
| CD138 Biotin | 1:1000 | 142512 | Biolegend | AB_2561981 |
| IgG2b Biotin | 1:1000 | 406704 | Biolegend | AB_315067 |
| CD8α Biotin | 1:1000 | 100704 | Biolegend | AB_312743 |
| CD4 Biotin | 1:1000 | 100404 | Biolegend | AB_312689 |
| IgG1 Biotin | 1:1000 | 406604 | Biolegend | AB_315063 |
| IgMa Biotin | 1:100 | 408603 | Biolegend | AB_940535 |

| | | | | |
|---|---|---|---|---|
| IgMa Purified | 1:100 | 408602 | Biolegend | AB_940549 |
| IgMb Biotin | 1:100 | 406204 | Biolegend | AB_315037 |
| IgMb Purified | 1:100 | 406202 | Biolegend | AB_315035 |
| Goat anti-mouse IgA, unlabeled | 10 ug/ml | 1040-01 | Southern Biotech | AB_2314669 |
| Goat anti-mouse IgG2b, unlabeled | 10 ug/ml | 1090-01 | Southern Biotech | AB_2794517 |
| Goat anti-mouse IgM, unlabeled | 10 ug/ml | 1020-01 | Southern Biotech | AB_2794197 |
| BRDU FITC kit | | 559619 | BD Biosciences | AB_2617060 |
| Rabbit Anti-RFP | 1:800 | 600401379 | Rockland | AB_11182807 |
| DAPI | 1:4500 | | | |
| GL7 bio for section | 1:500 | 144616 | Biolegend | AB_2721505 |
| CD21/35 for section AF647 | 1:100 | | | |
| Ibrutinib | | | | |
| R406 | | | | |
| BD Cytofix/Cytoperm | | 554714 | BD Biosciences | AB_2869008 |
| Cholera Toxin | | C8052-2MG | Sigma | |
| Fluo 3 AM | | F23915 | Invitrogen | |
| Fura Red | | F3021 | Invitrogen | |
| p-Tyrosine mouse HRP | 1:100 | 5465S | Cell Signaling | |
| 16% PFA | | 15710 | Electrom microscopy sciences | |
| 10% Mini-PROTEAN® TGX™ Precast Protein Gels | | 4561033 | Biorad | |
| SuperSignal West Pico Plus substrate | | 34577 | Thermo Fisher | |
| EasySep Streptavidin RapidSpheres Kit | | 19860A | Stemcell Technologies | |
| Trans-Blot Turbo Mini 0.2 µm PVDF Transfer Packs | | 1704156 | BioRad | |
| Protein Standard | | 161-0374 | BioRad | |
| ECL prime western blotting detection | | RPN2232 | GE Healthcare | |
| Anti-Phosphotyrosine | 1:2000 | 05-321 | Millipore Sigma | AB_309678 |
| Anti β-Actin | 1:5000 | A5316 | Millipore Sigma | AB_476743 |
| Anti-mouse IgG-HRP | 1:5000 | 7074S | Cell Signaling Technology | |

### Supplementary Figure Legends

#### Supplemental Figure 1

(A) Immunohistochemisty ratio of GC to follicle size of *Iga*$^{+/+}$ and *Iga*$^{-/-}$ Peyer's patches. (B) Representative flow cytometry of Peyer's patches. Gated previously on live singlets. (C) CD73- CD80- (DN) MemB as percentage of activated (IgD-) B cells in *Iga*$^{+/+}$ and *Iga*$^{-/-}$ mice. (D) mLN Foll as percentage of B cells. GC, memory B, and PCs as percentage of activated B cells from *Iga*$^{+/+}$ and *Iga*$^{-/-}$ mice. (E, F) Isotypes of GC B cells in PPs(E) and mLN (F). (G, H) Isotypes of memory B cells from PPs(G) and mLN(H). (I, J) Isotypes of PCs from PPs(I) and mLN(J). A-J, compiled data from at least 3 independent experiments with 1-3 mice per group. (K) Representative serial IHC sections of mBM chimeras stained with IgD (brown) and CD45.1 or CD45.2 (blue). Scale bars, 100 um. (L) Ratio of frequency of CD45.2 DN memB cells to CD45.2 Foll B cells in PP and mLN of Salmonella infected mBM chimeras. Unpaired t-test was used in a, c, d, e, f, g, h, i, j, l. ns=not significant, *p<0.005, **p<0.001,***p<0.0005, ****p<0.0001.

#### Supplemental Figure 2

(A) Isotypes of gut homing ($\alpha 4\beta 7$+ CCR9+) and non-gut homing PCs from mLN of WT mice treated with FTY-720 for seven days. Compiled data from 2 independent experiments with 2-3 mice per group.
(B) Representative immunofluorescence of Peyer's patches and small intestinal villi of *Aidca*$^{cre/+}$ R26$^{tdTomato}$ *Iga*$^{+/+}$, *Aidca*$^{cre/+}$ R26$^{tdTomato}$ *Iga*$^{+/+}$, *Aidca*$^{cre/cre}$ R26$^{tdTomato}$ mice stained with DAPI (gray) and tdTomato (red). Scale bars, 200 um. (C) Ratio of frequency of CD45.2 PCs to CD45.2 Foll B cells in PP and mLN of mBM chimeras. Representative data from at least 3 independent experiments with at least 3 mice per group. (D) Representative immunofluorescence of small intestinal villi stained for DAPI (gray), tdTomato (red), and IgA (yellow), in WT: *Aidca*$^{cre}$ R26$^{tdTomato}$ *Iga*$^{+/+}$ mBM chimera. Scale bars, 50 um. Inserts of IgA+ tdTomato+ cells within villi, scale bar 10 uM. (E) Ratio of frequency of IgM$^{a}$ to IgM$^{b}$ in indicated B cell populations from PP and mLN of WT and *Iga*$^{+/-}$ mice. Compiled data from 4 independent experiments with 2-4 mice per group. (F) Ratio of frequency of IgM+ CD45.2 B cells to IgM+ CD45.1 for indicated B cell populations of PP and mLN of mBM chimeras. Representative data from at least 3 independent experiments with at least 3 mice per group. (G) Percentage of endogenously coated fecal bacteria with indicated antibodies. One way anova was used in a. Unpaired t-test was used in c, e, f, g. ns=not significant, *p<0.005, **p<0.001,***p<0.0005, ****p<0.0001.

#### Supplemental Figure 3

(A) Ratio of frequency of CD45.2 LZ or DZ GC B cells to CD45.2 Foll B cells in mLN of mBM chimeras. Data is representative of at least 3 independent experiments with at least 3 mice per group. (B) Representative histograms of BRDU staining in PP DZ and LZ GC B cells of mBM chimeras. (C) Percentage of BRDU+ DZ or LZ GC B cells in PPs of *Iga*$^{+/+}$ :WT mBM chimeras after 3.5 hour BRDU pulse. (D) Percentage of BRDU+ DZ and LZ GC B cells from mLN of indicated mBM chimeras. (E) Percentage of BRDU+ Foll B cells from PP or mLN of indicated mBM chimeras. C-E**,** Compiled data from at least 3 independent experiments with at least 3 mice per group. (F) Ratio of frequency of specified CD45.2 B cells to CD45.2 Foll B cells in mLN of BCL2 retroviral mBM chimeras. Compiled data from 2 experiments with at least 3 mice per group. (G) Representative active caspase 3 staining in DZ and LZ GC B cells of mBM chimeras. (H) Percentage of active caspase3+ DZ GC B cells from PPs of mBM chimeras. (I) Percentage of active caspase3+ DZ and LZ GC B cells from mLN of mBM chimeras. Compiled data from at least 3 independent experiments with at least 3 mice per group. (J) surface IA-IE gMFI on WT PP GC B cells for indicated isotypes. Compiled data from at least 3 independent experiments with at least 2 mice per group. (K) Percentage of BRDU+ LZ GC B cells in mLN for indicated isotypes after 30-minute BRDU pulse. (IgG are defined as IgM- IgA-). Compiled data from two experiments with 3 mice per group. Unpaired t-test was used in a, c, d, e, h, i. One-way anova was used in f, j, k. ns=not significant, *p<0.005, **p<0.001,***p<0.0005, ****p<0.0001.

#### Supplemental Figure 4

(A) PP GC surface gMFI for indicated BCR components on specified isotypes**.** Compiled data from at least 3 independent experiments with at least 2 mice per group. (B) Representative flow cytometry of *Aidca*$^{cre/+}$ R26$^{td/gCAMP}$ PP GC B cells stimulated with indicated anti-BCR. (C) Compiled calcium traces for IgM, IgG2b, and IgA memory B cells (defined as isotype negative populations) stimulated with anti-panBCR. (D) Area under the curve of individual calcium traces calculated between 50 and 100 seconds for C. C, D are compiled data from

4 independent experiments. (E) Representative flow cytometry of *Aidca*$^{cre/+}$ R26$^{td/gCAMP}$ PP memory B cells stimulated with indicated anti-BCR. (F) Compiled calcium traces for *Aidca*$^{cre/+}$ R26$^{td/+}$ PP memory B cells stimulated with indicated anti-BCR isotypes. (G) Area under the curve of individual calcium traces calculated between 40 and 100 seconds for F**.** Indicated samples were pre-treated with ibrutinib for 10 minutes prior to anti-BCR stimulation. F, G are compiled data from at least 3 independent experiments. (H) Compiled calcium traces from *Aidca*$^{cre/+}$ R26$^{td/gCAMP}$ PP memory B cells stimulated with indicated anti-BCR isotypes. Foll B cell stimulated with anti-IgM. Shown as ratio of frequency of gCAMP-GFP+ to gCAMP-GFP- cells per second. (I) Area under the curve of individual calcium traces calculated between 50 and 100 seconds for H. H, I are compiled data from 4 independent experiments. J, K are compiled calcium traces for *Aidca*$^{cre/+}$ R26$^{td/+}$ PP GC(J) or memory B cells(K) stimulated with indicated anti-IgA. Indicated samples were pre-treated with ibrutinib for 10 minutes prior to anti-IgA stimulation. L, M are compiled calcium traces from *Aidca*$^{cre/+}$ R26$^{td/gCAMP}$ GC(L) or memory B cells(M) stimulated with anti-IgA. Shown as ratio of frequency of gCAMP-GFP+ to gCAMP-GFP- cells per second. Indicated samples were pre-treated with ibrutinib for 10 minutes prior to anti-BCR stimulation. One-way anova was used in a, c, g, i. ns=not significant, *p<0.005, **p<0.001,***p<0.0005, ****p<0.0001.

**Supplemental Figure 5**

(A) Representative flow cytometry of CH12 cell line after in vitro class switch to IgA. CH12 expressing surface IgM or IgA were sorted to produce uniform cell lines expressing IgM or IgA BCR. (B) Representative flow cytometry of I29 cell line in vitro class switch to IgA. I29 expressing surface IgM or IgA were sorted to produce uniform cell lines expressing IgM or IgA BCR. (C) pY gMFI normalized to untreated in CH12 cells after anti-BCR stimulation. (D) Representative pBTK histogram in CH12 cells in after anti-BCR. (E) pY, pBTK, pSyk gMFI normalized to untreated in I29 cells after anti-BCR stimulation. Compiled from at least 3 independent experiments. (F) pY gMFI in unstimulated CH12 cells. (G) pY gMFI in unstimulated I29 cells. (C, F, G) are compiled from 2 independent experiments. (H) Representative histogram of pBTK after incubation with R406 in CH12 cells. (I) pY or pSyk gMFI normalized to untreated in CH12 cells after incubation with R406. Compiled data from at least 3 independent experiments. (J) pY, pBTK, pSyk gMFI normalized to untreated in I29 cells after incubation with R406. Compiled data from at least 3 independent experiments. Unpaired t-test was used in c, e, f, g, I, j. ns=not significant, *p<0.005, **p<0.001,***p<0.0005, ****p<0.0001.

**Supplemental Figure 6**

(A) Representative histograms of surface Fas on CH12 cells. (B) Surface Fas gMFI on PP GC B cells for indicated isotypes. Compiled data from at least 3 independent experiments with at least 2 mice per group. (C) Ratio of frequency of CD45.2 LZ or DZ B cells to CD45.2 Foll B cells in mLN of mBM chimeras. (D) Ratio of frequency of CD45.2 CD73+ CD80+ memory B cells to CD45.1 memory B cells from mLN of mBM chimeras. (E, F) Isotypes of PP(E) and mLN(F) for indicated B cells from mBM chimeras. (G) Ratio of frequency of CD45.2 PCs to CD45.1 PCs from PPs and mLN of mBM chimeras. C-G are compiled data from 3 independent experiments with 1-3 mice per group. (H) Percentage of endogenously coated small intestinal bacteria with IgM$^a$ or IgM$^b$ antibodies from serum of mBM chimeras made in uMT hosts. Percentage shown as IgM$^a$/IgM$^b$ ratio. Compiled data from 3 independent experiments with 2-3 mice per group. One-way anova was used in b-h. ns=not significant, *p<0.005, **p<0.001,***p<0.0005, ****p<0.0001.

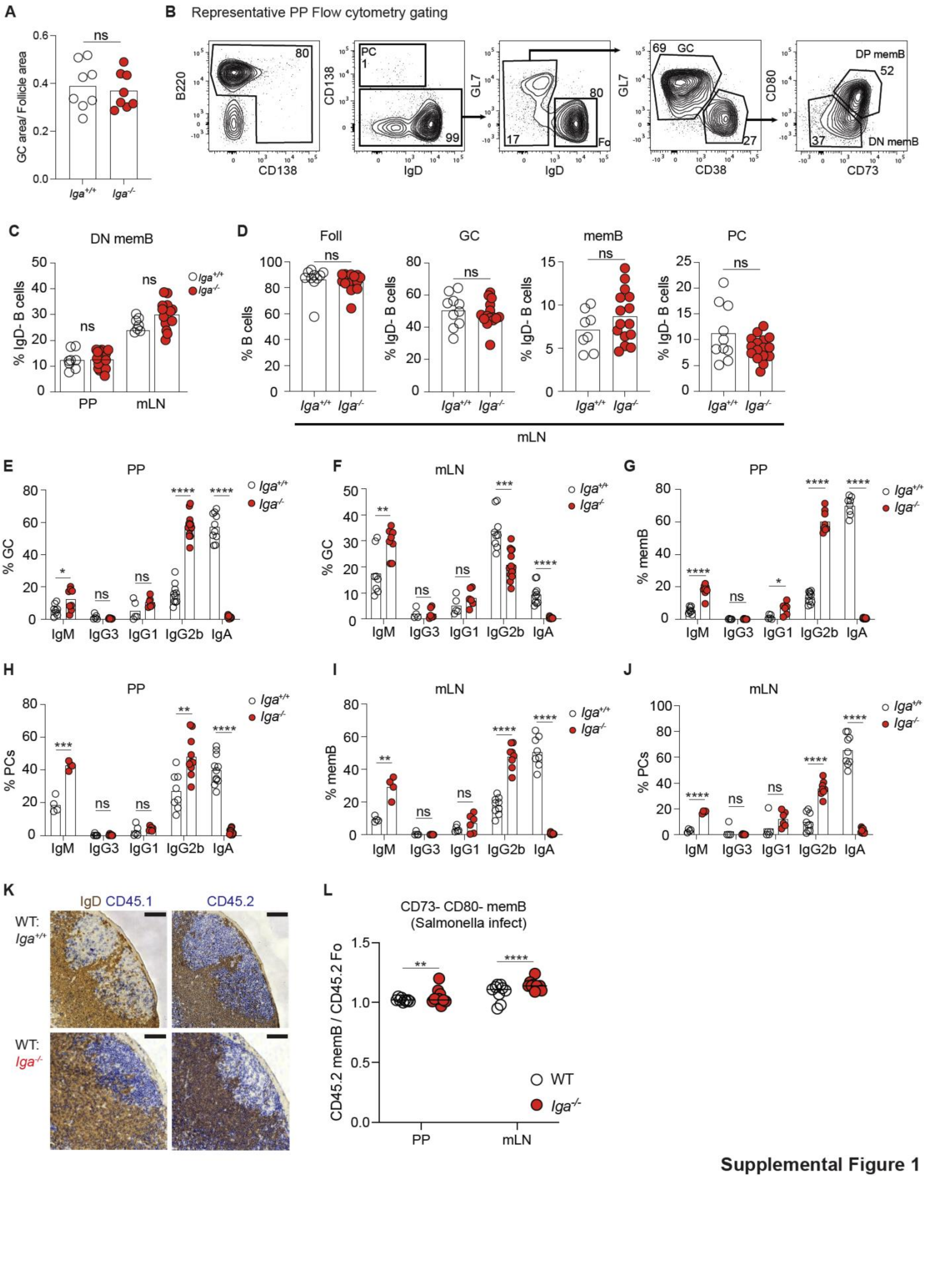


Supplemental Figure 1

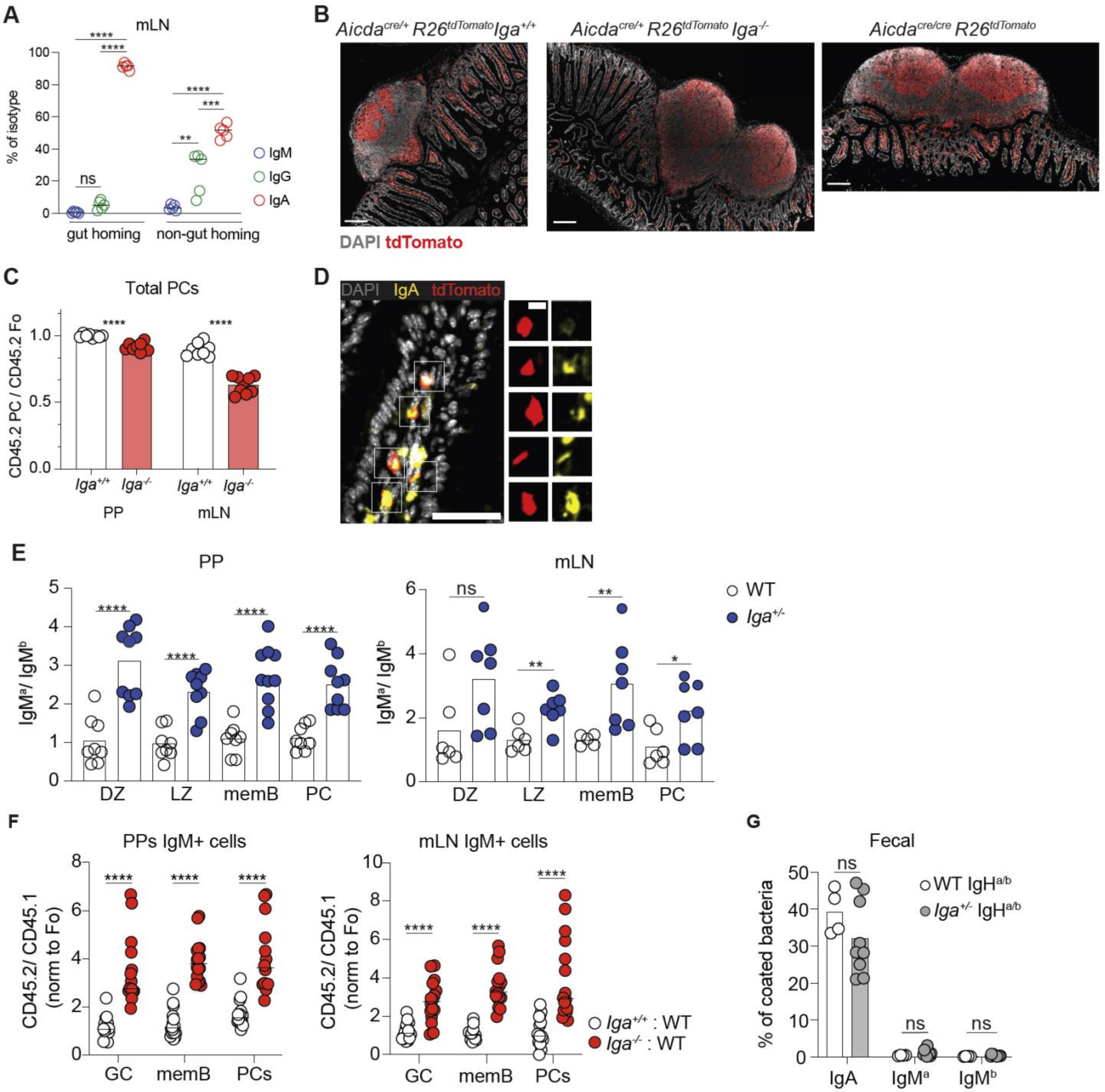


**Supplemental Figure 2**

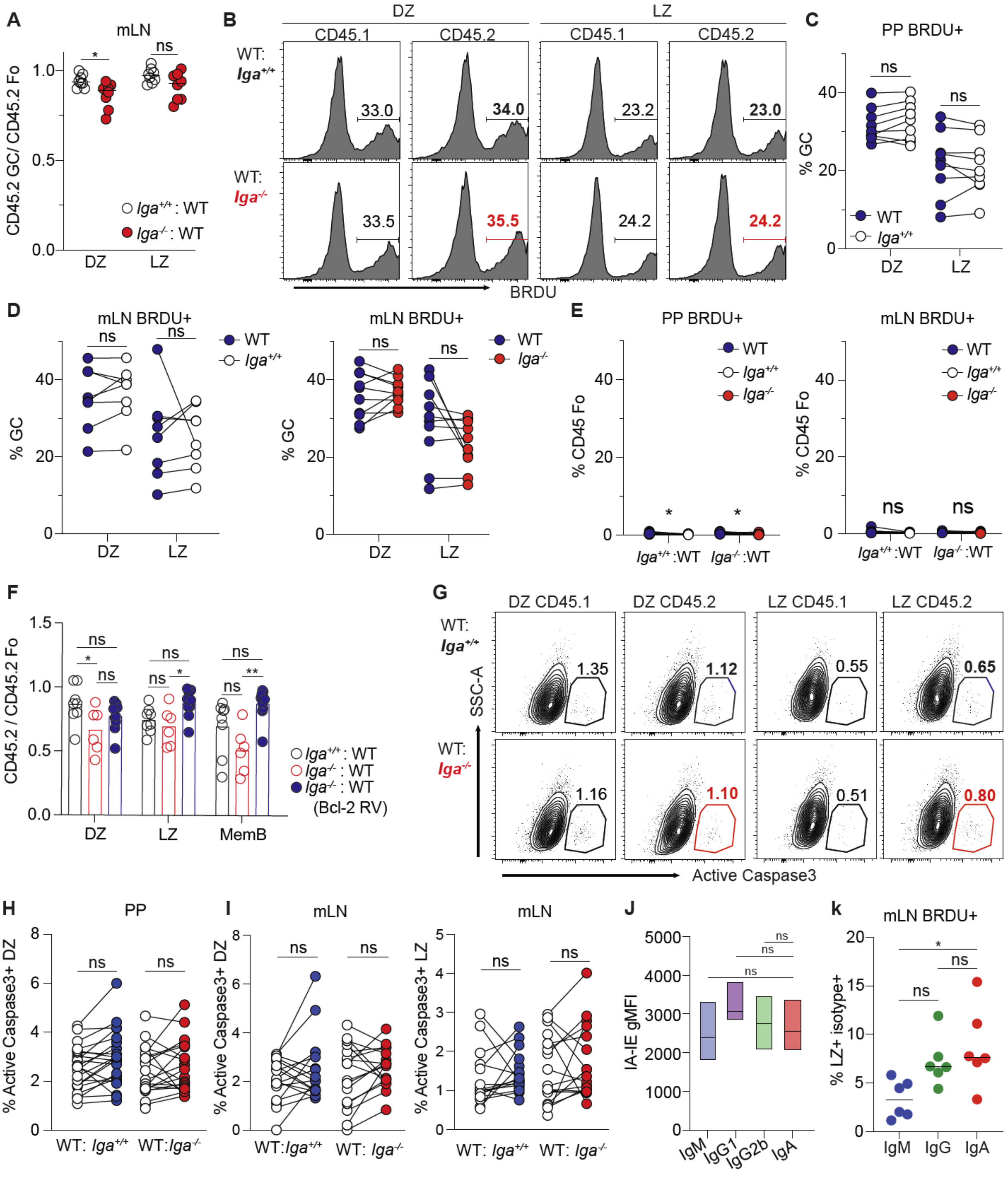


**Supplemental Figure 3**

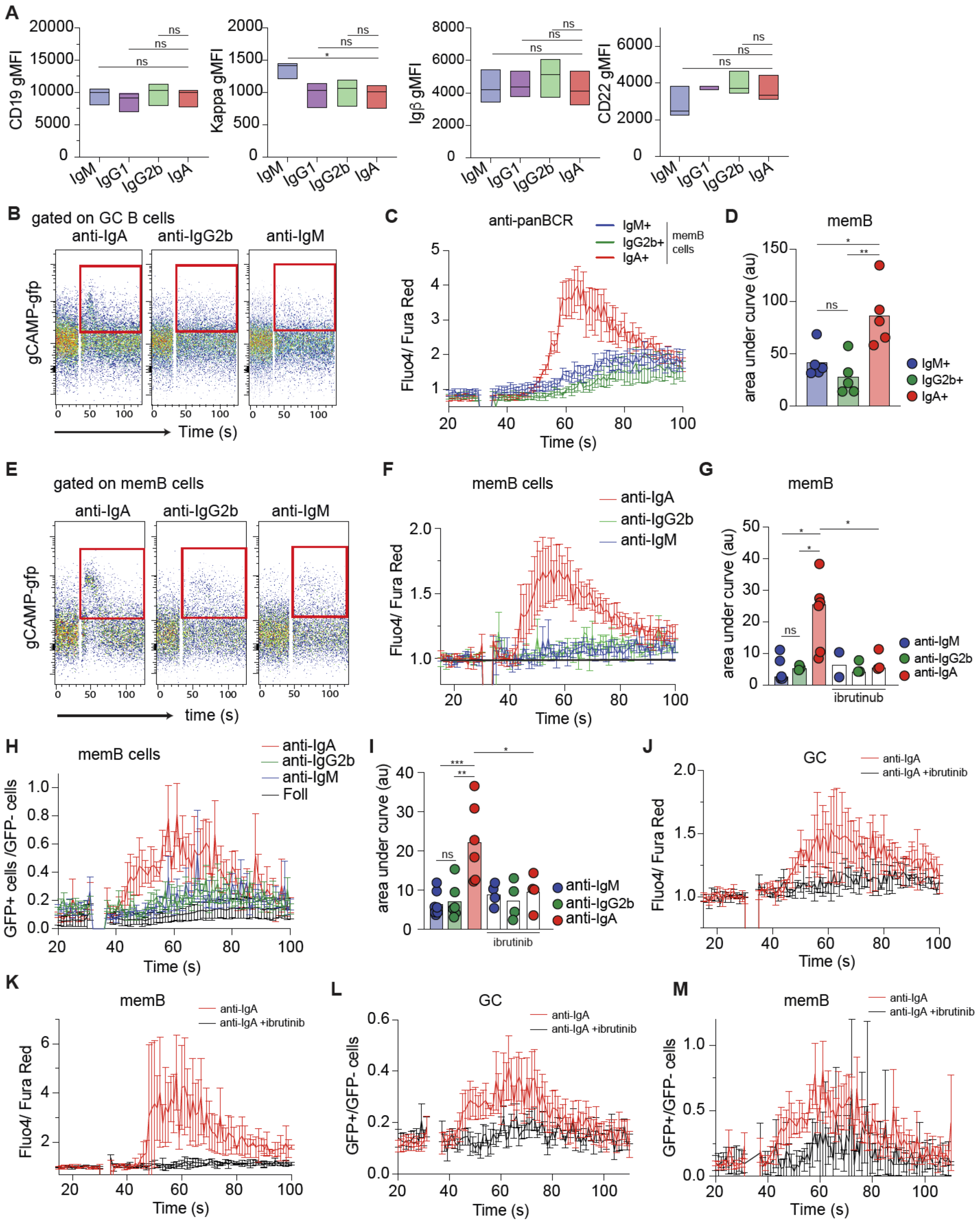


**Supplemental Figure 4**

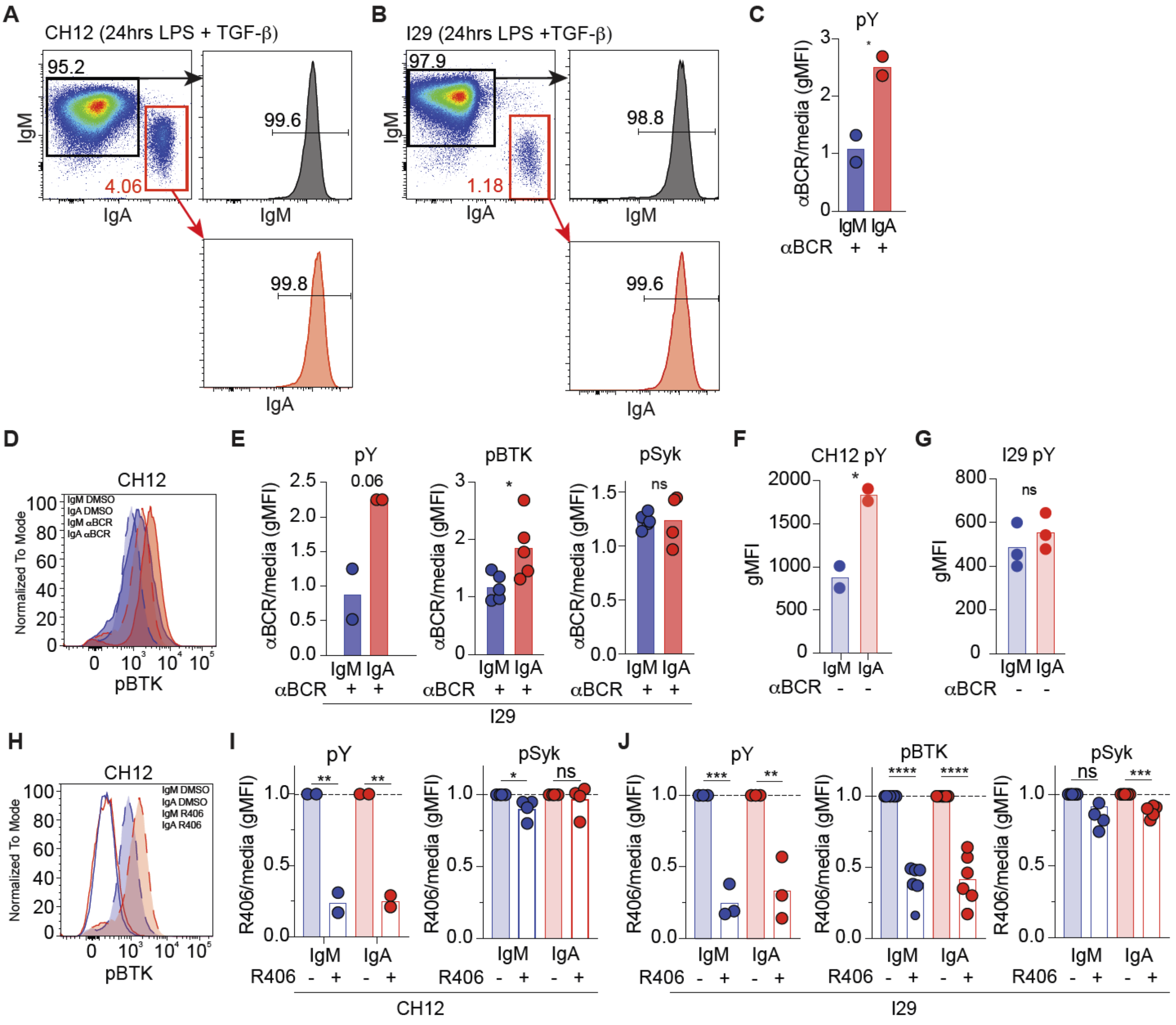


**Supplemental Figure 5**

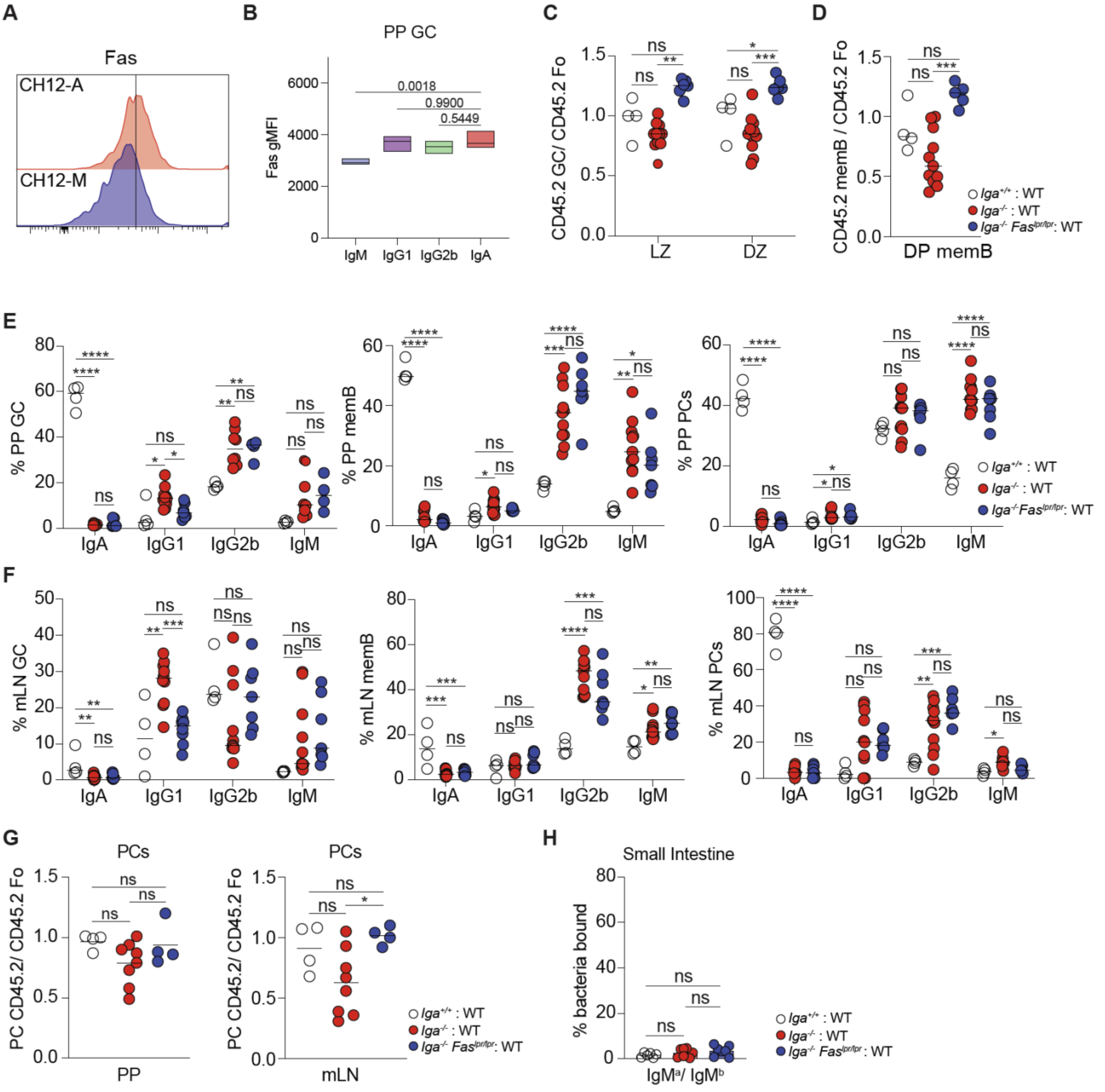


**Supplemental Figure 6**
